# Gene regulatory network inference from single-cell data using multivariate information measures

**DOI:** 10.1101/082099

**Authors:** Thalia E. Chan, Michael P.H. Stumpf, Ann C. Babtie

## Abstract

While single-cell gene expression experiments present new challenges for data processing, the cell-to-cell variability observed also reveals statistical relationships that can be used by information theory. Here, we use multivariate information theory to explore the statistical dependencies between triplets of genes in single-cell gene expression datasets. We develop PIDC, a fast, efficient algorithm that uses partial information decomposition (PID) to identify regulatory relationships between genes. We thoroughly evaluate the performance of our algorithm and demonstrate that the higher order information captured by PIDC allows it to outperform pairwise mutual information-based algorithms when recovering true relationships present in simulated data. We also infer gene regulatory networks from three experimental single-cell data sets and illustrate how network context, choices made during analysis, and sources of variability affect network inference. PIDC tutorials and open-source software for estimating PID are available here: https://github.com/Tchanders/network_inference_tutorials. PIDC should facilitate the identification of putative functional relationships and mechanistic hypotheses from single-cell transcriptomic data.

## Introduction

Precisely controlled patterns of gene expression are essential for the survival and reproduction of all life-forms. Development provides the canonical example, where changes in gene regulation determine the path by which from a single fertilised egg cell emerges a complex multicellular organism. Intricate networks of transcriptional activators and repressors have evolved to regulate the spatial and temporal expression of genes, enabling organisms to adjust transcription levels in response to environmental, developmental and physiological cues (Trapnell et al. 2014, Harrington et al. 2014, Rué et al. 2015, Moris et al. 2016, Gouti et al. 2015, Göttgens 2015). Elucidating the structure of such gene-regulatory networks (GRNs) has been a central goal of much recent systems biology research (De Smet et al. 2010, Oates et al. 2012, Thorne et al. 2012, Thorne et al. 2013, Siegenthaler et al. 2014, Oates et al. 2014, Huang et al. 2014, Young et al. 2014), and it is now becoming a pivotal stepping stone in dissecting the molecular contributions of complex diseases (Boyle et al. 2017).

The structure of GRNs alone does not fully constrain their function (Ingram et al. 2006), but it serves as an important starting point for further analysis. The simplest mathematical representations of GRNs are static, undirected graphs, where each node represents a gene, and edges depict relationships between transcription factors and their targets. Although by this definition the GRN encapsulates every transcriptional regulatory relationship that could occur within a given organism, this is not a very helpful perspective: clearly, it is not the case that every possible interaction does occur in every cell — for example, the downstream interactions of a transcription factor only occur when it is expressed — hence we can define subsets of the GRN that are active in particular cells and contexts. The structure and dynamics of these active GRN subsets give rise to distinct mRNA expression profiles, and it has been suggested that characteristic expression profiles in different cell types (and under different conditions) result from different stable states of the GRN (Clevers et al. 2017, Huang 2010, Moris et al. 2016, Moignard et al. 2015).

The introduction of efficient high-throughput expression assays has driven interest in *network inference* methods that apply statistical approaches to identify likely regulatory relationships between genes based on their expression patterns and potential GRN structures. In addition to correlation-based networks (perhaps the simplest way of identifying putative relationships), Gaussian graphical models, (Dynamical) Bayesian networks, regression analyses, and information theoretical approaches have been used for network inference from population level data (Penfold et al. 2011, Penfold et al. 2015, Bonneau et al. 2006, Margolin et al. 2006a, Villaverde et al. 2013, Villaverde et al. 2013, Liang et al. 2008, Madar et al. 2010, Hill et al. 2012, Lèbre et al. 2010, Beal et al. 2005, Schäfer et al. 2005, Vinciotti et al. 2016). Combining multiple inferred networks to form a community or ensemble prediction often confers slight but consistent improvements in the quality of the predicted network (Hill et al. 2016, Marbach et al. 2010, Marbach et al. 2012), but how to best combine and weight different methods to form a consensus prediction is poorly understood. Given our current understanding, a reasonable approach to generate such ensemble predictions would be to include information derived using different classes of inference algorithms (since these are known to show different biases (Marbach et al. 2012)) but also to ensure that within each class we develop the best performing algorithm based on a given statistical methodology.

More recently, the increasing availability of single cell expression data has led to the development of several computational and statistical approaches aimed at gaining new insight into cell fate decisions and transitions between cell states (Pina et al. 2015, Moignard et al. 2015, Rué & Martinez Arias 2015, Moris et al. 2016, Bendall et al. 2014, Trapnell et al. 2014). Identifying the associated changes in transcriptional state, and regulatory interactions that contribute to controlling these processes are key aims of many single-cell transcriptomic studies (Kharchenko et al. 2014, Ocone et al. 2015, Moignard et al. 2015, Pina et al. 2015, Trapnell et al. 2014, Bendall et al. 2014). Several *pseudotemporal ordering* algorithms have been developed that aim to place cells in an inferred temporal order based on similarities in their transcriptional states (Trapnell et al. 2014, Bendall et al. 2014, Haghverdi et al. 2016, Reid et al. 2016, Setty et al. 2016), since true single-cell temporal data (where large numbers of genes are assayed) are not feasible to collect at present. These methods often make strong assumptions about developmental processes (that have been questioned (Moris et al. 2016)) and, in most cases, uncertainties in the inferred order are likely to affect and bias downstream analyses. Network inference methods, in contrast, explore statistical dependencies between genes (from the observed distributions of expression levels across a given population of cells) and identify those that may be indicative of functional relationships, without necessarily making such strong assumptions about the nature of cell transitions; each inferred edge is a hypotheses that can be tested. Information theoretical measures, in particular, are particularly parsimonious in the assumptions that they make (Kinney et al. 2014) compared to e.g. ODE-based regression approaches or simple graphical models. However, network inference using single cell data (Fig. 1) remains relatively unexplored with only a few notable examples, e.g. (Ocone et al. 2015, Moignard et al. 2015, Filippi et al. 2016), extending beyond simple, potentially simplistic, correlation-based analyses (Bacher et al. 2016, Moignard et al. 2013, Kolodziejczyk et al. 2015, Pina et al. 2015, Stegle et al. 2015), perhaps due to the difficulties of directly applying and interpreting the results of existing methods designed to deal with population-level data.

**Figure 1:**
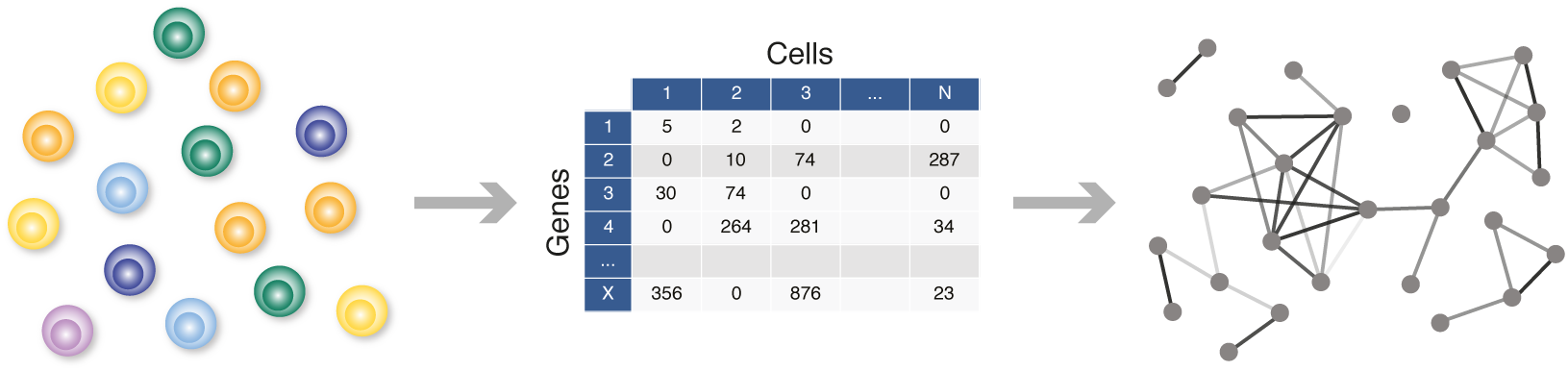
Network inference from single cell data. Single cell transcriptomic data quantify mRNA species present inside individual cells. By considering pairs (or triplets, quadruplets, etc.) of mRNA species we can test for statistical relationships among them. These dependencies may reflect coordinated gene expression of these pairs (or groups) of genes, resulting from gene regulatory interactions or co-regulation. Once such sets of genes that jointly change in expression are known, other statistical, bioinformatic, or text-mining analyses can be used to identify likely transcriptional regulators for these sets of genes. By iterating such in silico analyses with further, targeted experimental studies we can, in principle, build up a representation of the gene regulatory network.

Single cell data are notoriously complex and present new challenges for statistical analysis: technical noise is difficult to distinguish from genuine biological variability, the relative contributions and impact of different sources of noise are poorly understood, and numerous factors contribute to the biological heterogeneity observed within cell populations (Stegle et al. 2015, Pierson et al. 2015, Grün et al. 2015, Bacher & Kendziorski 2016, Liu et al. 2016, Buettner et al. 2015). However, these data also offer substantial advantages over population-level data that we can exploit in order to learn about the structure of GRNs governing the observed changes in gene expression. Firstly, datasets are large, routinely comprising expression measurements from hundreds or thousands of cells, and continuing advances in single cell technologies will allow further increases in sample sizes (Macosko et al. 2015, Klein et al. 2015). Additionally, single cell data inherently provide the variability required to detect statistical relationships between gene expression profiles (interpreted as putative functional relationships), whereas population-based studies need to introduce this variability by observing cell populations under different conditions, e.g. using time series or perturbation studies (Marbach et al. 2010, Marbach et al. 2012, Penfold & Wild 2011, Oates & Mukherjee 2012). While we can collect single cell time-series data, even data collected at one time point will contain variability due to i) asynchrony of cells within a population (in terms of progression through a biological process), and ii) biological heterogeneity and often the presence of multiple cell (sub)types.

Here, we introduce an information theory-based GRN inference algorithm designed to make use of these features of single cell data. Information theory provides a set of measures, chiefly among them *mutual information* (MI), that allow us to characterize statistical dependencies between pairs of random variables without making assumptions about the nature of the dependencies (Cover et al. 2012, Mc Mahon et al. 2014). MI has considerable advantages over simpler measures such as (Pearson) correlation, as it is capable of capturing complex non-linear and non-monotonic dependencies, and reflecting the dynamics between pairs or groups of genes (Mc Mahon et al. 2015, Uda et al. 2013). Calculating MI involves estimating pairwise joint probability distributions, generally requiring density estimation (Kraskov et al. 2004, Steuer et al. 2002) or data discretization, and the accuracy of these estimates depends on the sample sizes. Single cell datasets are sufficiently large to allow us to accurately estimate probability distributions between more than two variables, and thus capitalise on recent developments in multivariate information (MVI) theory. Quantifying the information between three or more variables has long been problematic, and the most widely-used measures, such as interaction information, are known to have serious flaws (Timme et al. 2014). The recently-introduced partial information decomposition (PID) both explains and solves its predecessors’ problems and provides a meaningful measure of MVI (Williams et al. 2010); PID and other measures are described in detail in Box 1. Our algorithm uses PID to analyse the statistical relationships between triplets of variables to generate undirected networks highlighting putative functional interactions between genes.

In this paper, we describe an inference algorithm based on the MVI measure PID and use extensive *in silico* analyses to demonstrate i) the consistent improvement over existing algorithms based on pairwise MI, and ii) the suitability of this method for analysing single cell data, before illustrating its application to several experimental datasets. Such *in silico* analyses are critical for quantitatively assessing network inference approaches as, unlike with real biological systems, we have knowledge of the ‘true’ GRN underlying the observed data. These results demonstrate that the larger sample sizes of single cell data are vital for our method and that they profoundly improve the performance of information-based methods in general. We thoroughly explore the factors that influence algorithm performance — in particular the choice of discretization algorithms and probability distribution estimators — in order to provide evidence-based guidelines for the use of information theoretical based methods for network inference. We emphasise the importance of considering the different sources of heterogeneity within single cell data so that we can take advantage of the variation of interest — e.g. that associated with progression through a biological process such as differentiation. Our examples using experimental data demonstrate how our inference method can be combined with existing computational and statistical methods (e.g. clustering and dimensionality reduction) to infer networks from carefully-chosen subsets of single cell data in order to address particular questions about cellular processes. We consider three single cell transcriptomic datasets here and additionally refer to a related manuscript (Stumpf et al. 2017), in which we use our framework to infer changing regulatory sub-networks over the course of neural progenitor development in mouse embryos, and to suggest candidate genes for maintaining cellular states and driving state transitions. Finally, we provide a fast, open-source implementation of our methods to enable easy application to other single cell datasets.

#### Box 1: Information theoretic measures

The entropy, *H*(*X*), quantifies the uncertainty in the probability distribution, *p*(*x*), of a random variable *X*. For a discrete random variable, the entropy is given by,

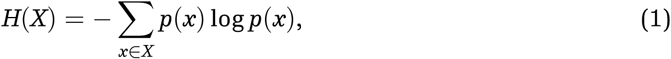

which is maximal for a uniform distribution. If we consider the mRNA expression level of a gene to be the variable *X*, then a gene that is expressed differently across a set of cells will have a higher entropy than a more consistently expressed gene. When considering the relationship between *X* and a second random variable, *Y*, we quantify the information that one variable provides about the other using the MI,

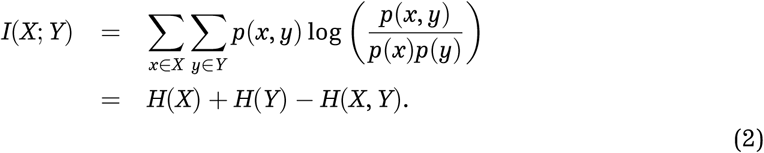

This quantifies the difference between the joint entropy, *H*(*X*, *Y*), and the joint entropy assuming independence of *X* and *Y*, *H*(*X*) + *H*(*Y*), and thus provides a non-negative, symmetric measure of the statistical dependency between the two variables. For a pair of genes with more co-ordinated expression, their observed joint entropy will be lower, and hence they will have higher mutual information. Given a third variable, *Z*, the conditional mutual information (CMI),

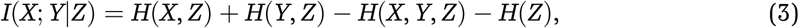

quantifies the information between *X* and *Y* given knowledge of *Z*. This tells us the extent to which knowing the expression of one gene additionally informs us about the expression of a second gene, given that we already know the expression of a third gene.

A number of MVI measures have been defined that aim to quantify the statistical dependencies between three or more variables, but there is little consensus as to the most appropriate metric (Timme et al. 2014). Arguably the most widely used is interaction information (McGill 1954), which for three genes quantifies the extra information between any two of the genes *X*, *Y* and *Z*, when the third is known compared to when it is not known,

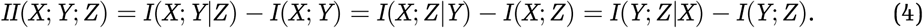

Interaction information has received much criticism since it can be (i) zero between dependent variables, when MI and CMI are equal but non-zero and (ii) negative, when MI is greater than CMI. The problem is that, despite its name, interaction information is not a quantity of information between a set of variables; rather it quantifies the balance between MI and CMI. In fact MVI is difficult to summarise with a single quantity, because there are different ways in which information can be shared by three or more variables.

An important recent development in information theory is the introduction of partial information decomposition (PID) (Williams & Beer 2010). In the three variable case, PID considers the information provided by a set of *source* variables (or genes), *S* = {*X*, *Y*}, about another *target* variable, *Z*, partitioned into redundant, synergistic and unique information. Redundant information is the portion of information about *Z* that can be provided by either variable in *S* alone; the unique contribution from *X* (or *Y*) is the portion of information provided only by *X* (or only *Y*); and the synergistic information is the portion of information that is only provided by knowledge of both *X* and *Y*. Thus the PID between the set *S* and the target variable *Z* is equal to the sum of the four partial information terms,

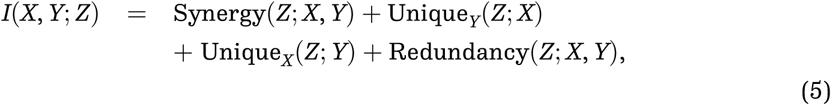

where Unique_*Y*_(*Z*; *X*) is the unique information between source variable *X* and target variable *Z* when the other source variable is *Y*. In the context of gene expression, the terms ‘source’ and ‘target’ do not imply any mechanistic assumptions; rather, they refer to quantifying to what extent knowledge of the source genes inform us about the target gene.

To calculate the PID terms, the redundant information is first calculated using the specific information, *I*_*spec*_, which quantifies the information provided by one variable about a specific state of another variable (Deweese et al. 1999, Timme et al. 2014). The ‘state’ of a gene in a given cell refers to which discrete bin its mRNA level falls within, once we have discretized the expression data. If we consider the information provided by *X* about state *z* of variable *Z*,

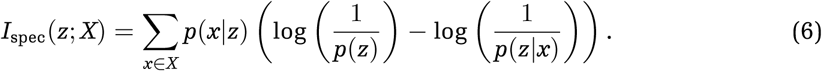

If we consider *S* = {*X*, *Y*} and a target variable *Z*, the redundant contribution is calculated by comparing the amount of information provided by each variable within *S* about each state of the target *Z*,

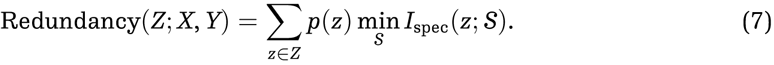

The unique information terms can be calculated from the redundant information and the pairwise MI, via the relationship,

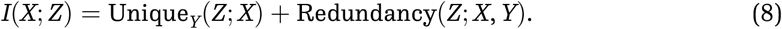

In other words, the pairwise MI between two variables can be partitioned into a redundant and unique component, given a third variable; it is this relationship that we exploit further in *Results*. We note that although PID is not a symmetric measure, Unique_*Y*_(*Z*; *X*)+Redundancy(*Z*; *X*, *Y*) will be equal to Unique_*Y*_(*X*; *Z*) + Redundancy(*X*; *Z*, *Y*) because MI is symmetric.

Finally the synergistic information can be calculated via the interaction information (Eq. 4), which turns out to be the difference between the synergistic and redundant information:

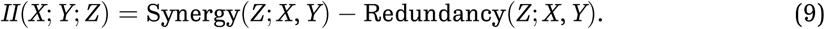

## Results

### PID profiles in synthetic data

We first investigate the usefulness of PID for inferring network edges using data generated from *in silico* models. We use stochastic simulations from simple directed 3-node networks of varying topologies, and estimate PID values (redundant, synergistic and unique information, defined in Box 1) from these simulated data. Simulations were generated from two model definitions, based on thermodynamic or mass action kinetics, both commonly used in systems biology to represent gene regulation, as described in *Methods*. A distinctive pattern is apparent in networks with a single directed edge between two genes (‘one edge’ topology, Fig. 3A) — the unique information between the two connected genes is notably higher than both the unique information between unconnected genes and the redundancy values between all three genes. With increasing numbers of edges within the network, this pattern is lost; this makes sense intuitively as with higher connectivity we expect to see increased synergistic or redundant contributions (Fig. 3A and S1). Note that the pattern can only be observed under simulation conditions that generate variability in the observed variables (i.e. statistical relationships are not detectable when the system is at steady state; Fig. S1).

**Figure 2:**
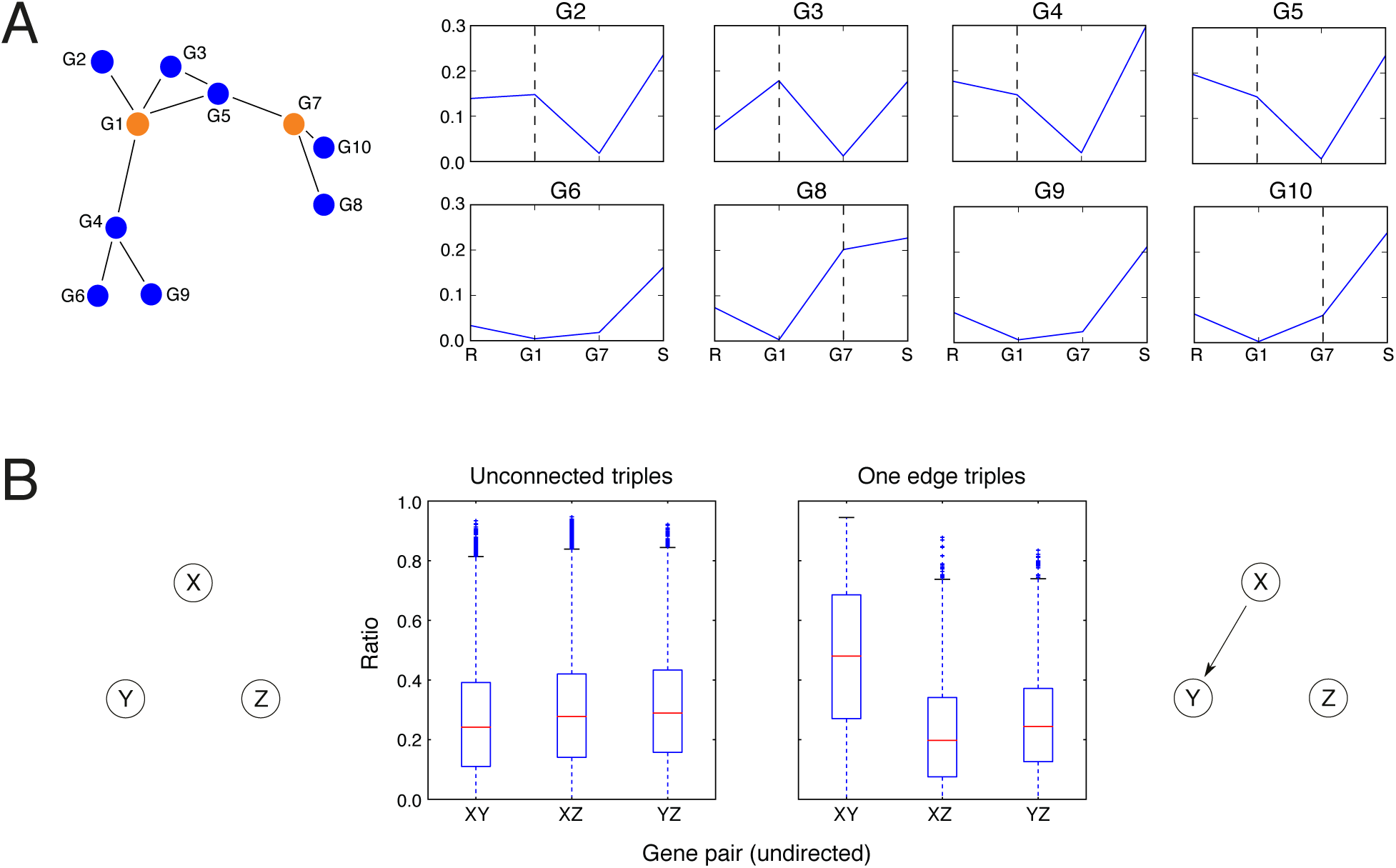
Demonstration of unique information in in silico networks. **A)** PID values are estimated from data simulated from a 10-gene in silico network (top) using `GeneNetWeaver` (Schaffter et al. 2011). Each line graph shows PID values estimated using genes 1 and 7 as the sources, and each of the remaining genes in turn as the target (graph titles, GX, indicate the target gene). Four PID values are given in each graph — the redundancy (R), the unique information between gene 1 and the target (G1), the unique information between gene 7 and the target (G7), and the synergy (S). The mutual information between two genes is the sum of their unique information and the redundancy (Eq. 8). The ratio of the unique information to the mutual information tends to be higher between pairs of connected genes (dashed vertical lines indicate the unique contributions for connected genes). **B)** Ratio of unique information to mutual information in triples with the two most common topologies, within the 50-gene network, S. cerevisiae 1. For each gene triple of the ‘unconnected’ and ‘one edge’ topologies (see Table S1 for topology frequencies), we calculate the unique information values between each pair of genes and their mutual information. The ratios of unique information to mutual information are higher in general for the connected pair; the same pattern was observed in all networks.

**Figure 3:**
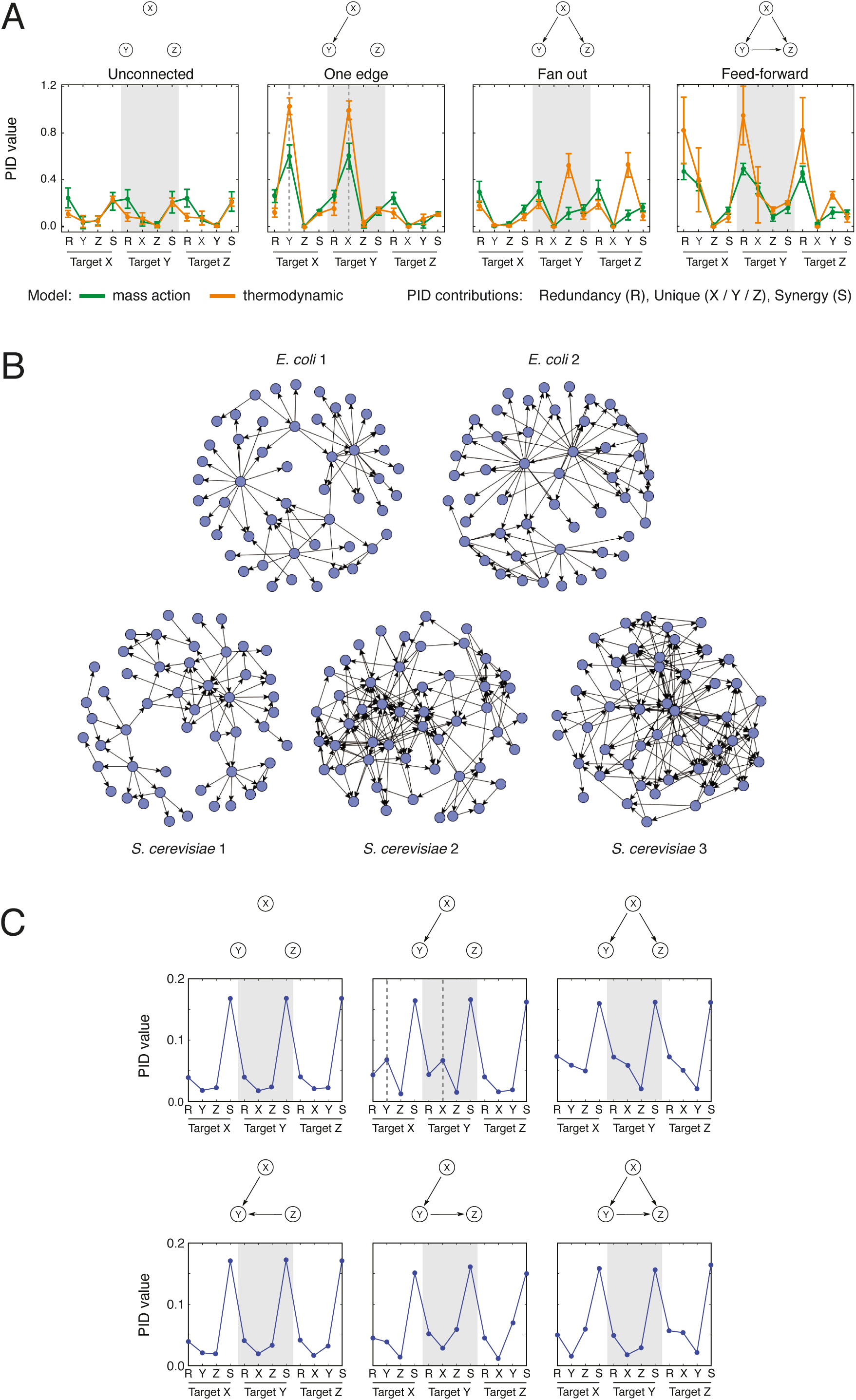
PID profiles for 3-gene networks. **A)** Mean PID values for 3-gene networks with different topologies. PID values are calculated using data simulated from 3-gene networks with the topologies illustrated above each plot; the models used for simulation assumed mass action (green) or thermodynamic (orange) kinetics. For each 3-gene network, twelve PID values were calculated from the simulated data — there are four PID contributions with each gene treated as the target gene in turn, consisting of a redundant, synergistic and two unique contributions (Eq. 4). Each line graph shows the mean PID values calculated from simulations using different initial conditions (error bars indicate 1 s.d.), with the horizontal axis labels indicating the PID contribution, e.g. the first four values show the PID values with gene X as the target, consisting of the redundancy (R), unique contributions from gene Y (Y) and gene Z (Z), and the synergistic contribution (S). The vertical dashed grey lines in the ‘one edge’ plot indicate the unique PID values that are used as the basis for our inference algorithm. Here, all regulatory interactions are assumed to be activating, the additional stimulating ligand targeted gene X, and the values indicated are the mean PID values calculated from five sets of simulations (with different randomly sampled initial conditions); results obtained with models that include both activating and inhibitory regulation are shown in Fig. S1. **B)** Visualisations of the directed 50-node networks, produced by GeneNetWeaver (Schaffter et al. 2011); node degree distributions for the 50- and 100-node GeneNetWeaver networks used in this study are shown in Fig. S2. **C)** Mean PID profiles for gene triplets in the 50-gene S. cerevisiae 1 network. Every triplet of nodes (genes) in the network was assigned to one of six possible classes (based on the known connectivity of genes, as indicated in network diagrams above each plot). Each line graph shows the mean PID values calculated across triplets with the same topology, with the horizontal axis labels indicating the PID contribution.

To explore whether this pattern also occurs for triplets of nodes embedded in large networks, we consider time series expression data simulated from five different 50-gene networks generated by GeneNetWeaver (Schaffter et al. 2011). This software generates stochastic simulations from dynamical models that represent transcription and translation using a thermodynamic approach, with network structures that are inspired by known gene connectivity patterns in *E.coli* and *S.cerevisiae* (Fig. 3B), and it has become a standard tool for performance evaluation of network inference algorithms (Schaffter et al. 2011, Marbach et al. 2010, Marbach et al. 2012). PID values are estimated from these data for every triplet of genes within the networks, and each triplet is classified according to its topology — six topological arrangements are possible given the model assumptions (maximum of one edge between each pair of nodes, no self-regulation, and no feedback loops). Mean PID values are calculated for each group, and the same distinctive pattern (high unique versus redundant contributions for connected genes) is apparent for triplets with a single directed edge (Fig. 3C). As with the 3-node simulations, the pattern is lost in topologies with more connections; however, the ‘unconnected’ and ‘one edge’ topologies are by far the most prevalent in all networks — comprising 64.4-93.2% and 6.3-29.7% of all triplets respectively, and jointly comprising over 90% of triplets in every network (Table S1 and Fig S2).

Examining *in silico* data from a 10-gene network suggests that the relative size of the unique information compared to the redundancy — i.e. the proportion of MI accounted for by the unique contribution (Eq. 8) — is more informative than the absolute unique information (Fig. 2A). Using data from the 50-node networks, we confirm that this is the case more generally. We take all triplets of the ‘unconnected’ and ‘one edge’ topologies (with each gene being able to take part in multiple triplets); then for each triplet, treating each gene in turn as the target, we estimate (i) the redundant information between all three genes, (ii) the unique information between one source gene and the target and (iii) the unique information between the second source gene and the target. We plot the ratio of unique information to MI (the sum of unique and redundant information) for each pair of genes in each triplet and find that this ratio is higher in general between connected pairs (Fig. 2B).

### Incorporating PID into an inference algorithm

In a network of *n* genes, given a pair of genes *X* and *Y*, there are *n* − 2 gene triplets involving the pair. The MI between *X* and *Y*, *I*(*X*; *Y*), is unaffected by the choice of the third gene, *Z*, because MI is a pairwise measure, but the unique information between *X* and *Y*, Unique_*Z*_(*X*; *Y*), varies depending on *Z*. Furthermore, the difference between *I*(*X*; *Y*) and Unique_*Z*_(*X*; *Y*) is equal to the redundancy between all three genes (Eq. 8), meaning that we can regard the ratio Unique_*Z*_(*X*; *Y*)/*I*(*X*; *Y*) as capturing the proportion of MI that is accounted for by unique information between *X* and *Y*, as opposed to redundant information between all three genes. We note that the vast majority of all possible gene triplets in our *E.coli* and *S.cerevisiae* networks are of the ‘unconnected’ or ‘one edge’ topology (Table S1), and that the ratio Unique_*Z*_(*X*; *Y*): *I*(*X*; *Y*) is higher between connected pairs in a ‘one edge’ triple (Fig. 2). Therefore, we would expect that if *X* and *Y* are connected, then most of the triplets made with *X*, *Y* and all *n* – 2 possible *Z* in turn, would be of the ‘one edge’ topology — and likewise if *X* and *Y* are unconnected, then most triples involving *X* and *Y* would be of the ‘unconnected’ topology — so Unique_*Z*_(*X*; *Y*)/*I*(*X*; *Y*) would in general be higher if *X* and *Y* were connected.

We define the *proportional unique contribution* (PUC) between two genes *X* and *Y* as the sum of this ratio calculated using every other gene *Z* in a network (where *S* is the complete set of genes),

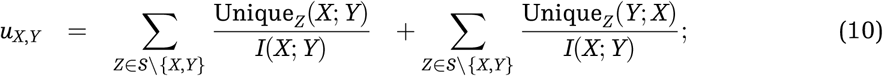

this measure may be thought of as capturing the mean proportion of MI between two genes *X* and *Y* that is accounted for by the unique information. Note that the PID unique measure is not symmetric, so for each pair of genes we consider each as the target in turn (hence we include both Unique_*Z*_(*X*; *Y*) and Unique_*Z*_(*Y*; *X*) terms in Eq. 10).

In our network inference algorithm (Fig. 4A), the redundancy and unique information contributions are first estimated for every gene triplet, then the PUC is calculated for each pair of genes in the network (Eq. 10). Finding a threshold for defining an edge at this stage is problematic, because the distribution of PUC scores differ between genes (see Fig. 4B), thus setting a global threshold for PUC scores across the whole network risks biasing the results by factors such as expression variability. This was previously observed with MI and led to the development of measures that take into account the network context, central to the *context likelihood of relatedness* (CLR) algorithm (Faith et al. 2007, Watkinson et al. 2009). A similar solution is employed here: an empirical probability distribution is estimated from the PUC scores for each gene, and the confidence of an edge between a pair of genes is given by,

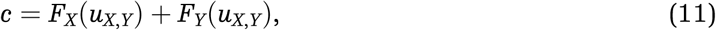

where *F*_*X*_(*U*) is the cumulative distribution function of all the PUC scores involving gene *X* (here, we assume either a Gamma or Gaussian empirical probability distribution). This effectively identifies the most important interactions per gene, rather than just taking the highest pairwise scores across the whole network.

**Figure 4:**
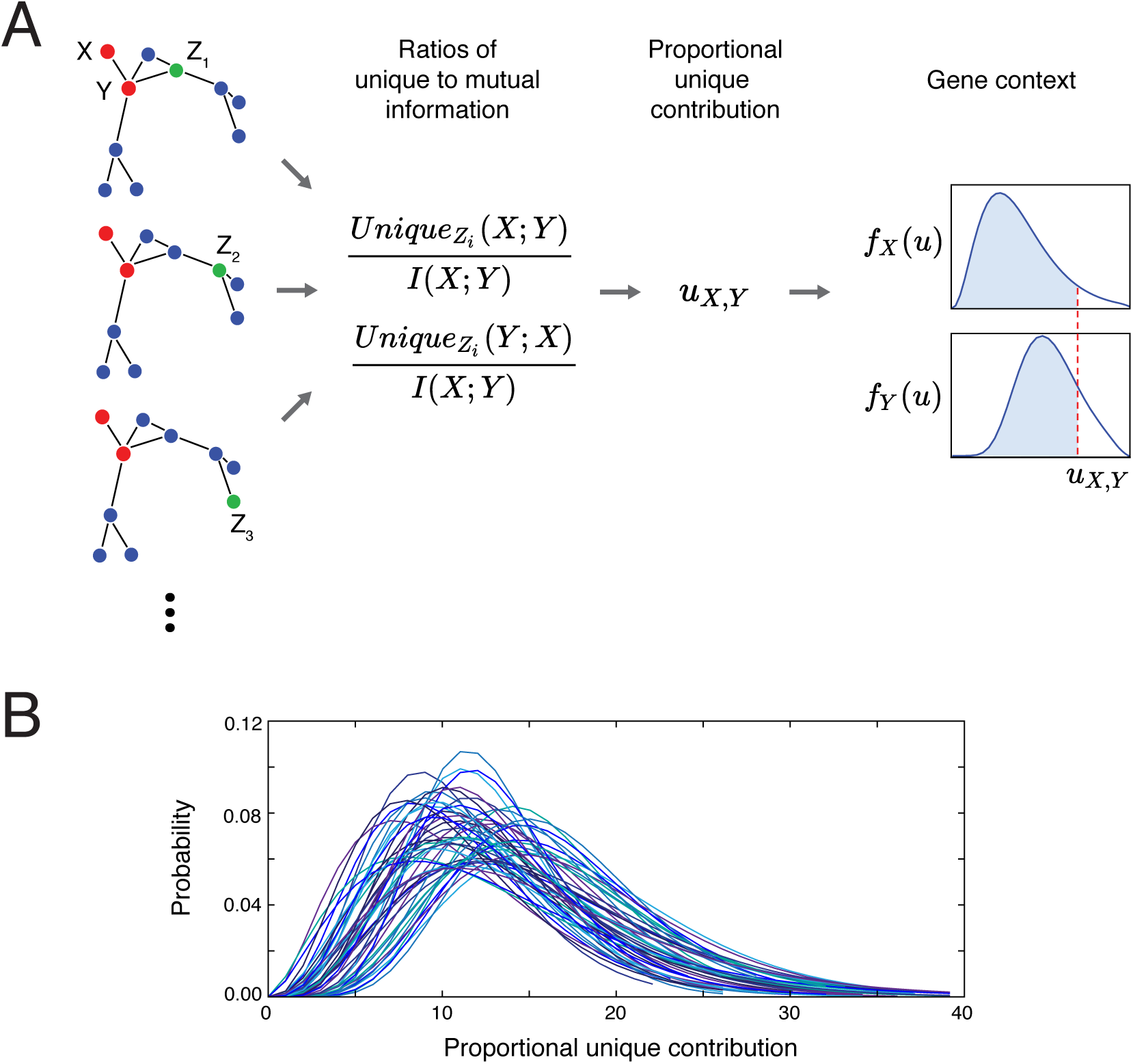
The PIDC inference algorithm. **A)** Schema of the PIDC inference algorithm. PID values are estimated for every gene triplet (with each gene treated as the target gene in turn), and from these the PUC, u_X,Y_, is estimated for every pair of genes. For each gene, X, an empirical distribution, f_X_(u), is estimated from its PUC scores with all other genes. The confidence of an edge between a pair of genes depends on the corresponding cumulative distribution functions, F_X_(u), for each gene within the pair (i.e. the blue shaded areas); these confidence scores are used to rank all possible network edges. **B)** Example empirical distributions of PUC scores by gene. Gamma distributions were fitted to the PUC scores (Eq. 7) for each gene in a 50-node in silico network (for each gene X, a PUC score, u_X,Y_, is obtained for that gene paired with each other gene Y in the network). Due to the variability of these distributions, using a universal threshold for inferring edges is problematic, thus we use the cumulative probability distributions for each gene to obtain a final confidence score for network edges. (Colours are to aid distinguishing the distributions.)

### Algorithm performance

We compare our algorithm against several common information theoretic based network inference methods, and thus briefly summarise these existing approaches in Box 2. All methods start from the pairwise MI matrix, and then use it in different ways. Even compiling the MI matrix is, however, fraught with potential problems: the manner in which data are treated (e.g. discretization), and the estimator used for the entropy and MI both affect the performance of the algorithms (Simoes et al. 2011, Zhang et al. 2015). When comparing different approaches it is therefore important to ensure that discretization and estimation of MI are done identically. Without this it becomes impossible to disentangle the relative strengths and weaknesses of the different approaches that are based on and interpret MI values. In the *Methods*, we discuss the different estimators and discretization approaches that we use and which are implemented in the InformationMeasures.jl package (see *Software*). Our comparisons with existing methods thus always start from the same MI matrix.

Undirected networks are inferred from *in silico* datasets (described in *Methods*) for five 50-gene net-works and five 100-gene networks using ARACNE, CLR, MI (relevance networks), MRNET (Meyer et al. 2008) and the PID-based algorithm, *PID and context* (PIDC). We also include results for the raw PUC score, without the network context step. Accuracy of the inferred networks is evaluated using the area under the precision-recall curve (AUPR), rather than the receiver operating characteristic curve (AUROC), which is inappropriate for judging network inference methods as real networks are typically sparse; see *Methods* for definitions and a more detailed discussion (Murphy 2012).

PIDC performs favourably compared to the other algorithms (Fig. 5), particularly in the larger networks. The raw PUC score outperforms the raw MI score, indicating the value of higher-order information; and CLR outperforms the other MI-based approaches, indicating the value of network context, in agreement with the previous comparisons (Fig. 5A and Fig. 5C) (Marbach et al. 2012). This effect is robust to simulated technical noise (Fig. 5B) and becomes more evident the larger the number of ‘cells’ in the dataset (Fig. 5A); as real data become more accurate and sample sizes increase, we expect the performance of PIDC/PUC to improve as the estimation of 3-dimensional dependencies becomes more accurate. Also, unlike CLR it was designed to capture higher order dependencies and to distinguish between direct and indirect interactions (Williams & Beer 2010).

**Figure 5:**
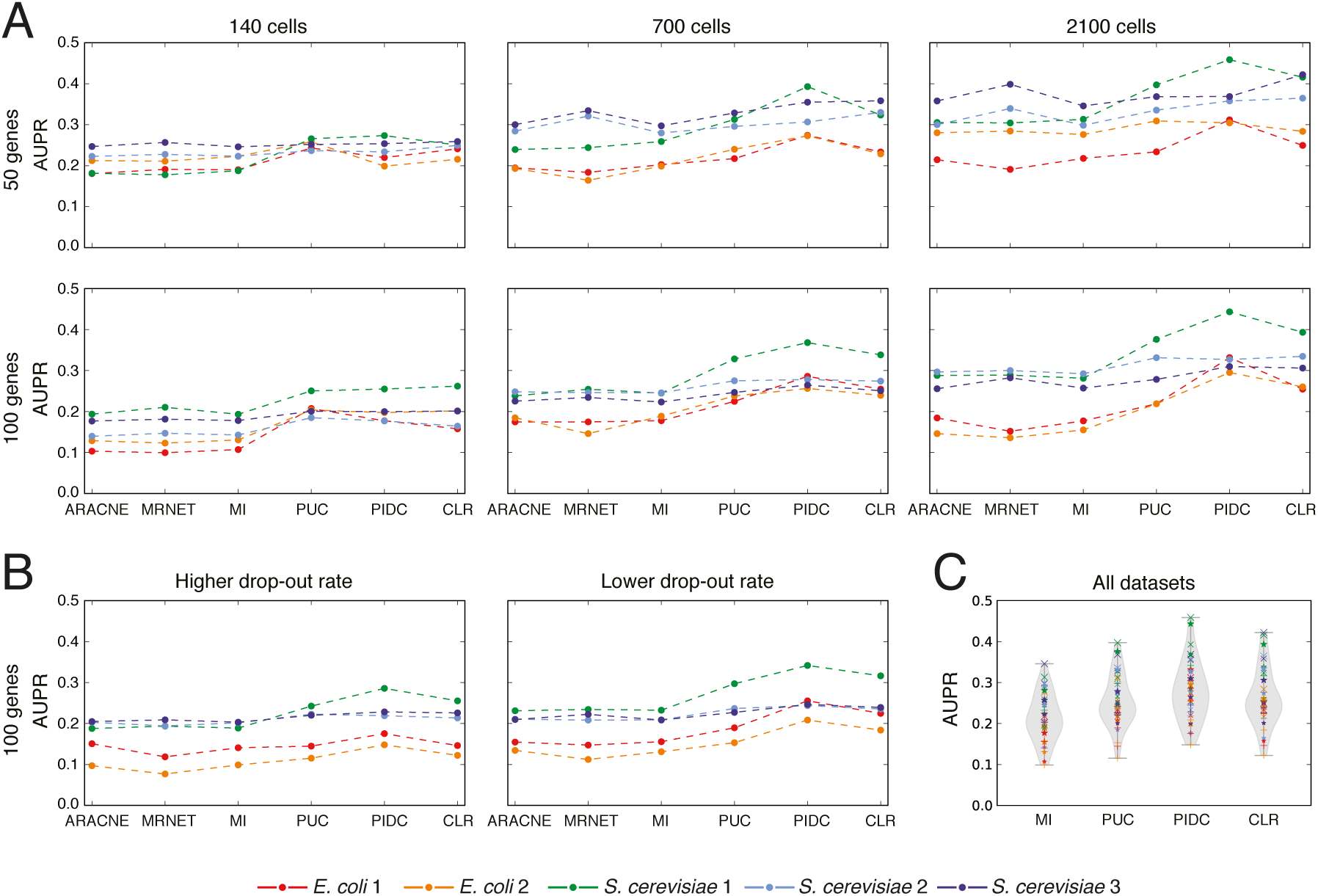
Performance comparison of information-theoretic network inference algorithms. **A)** AUPR is calculated for several algorithms applied to ten in silico datasets, generated from five 50-gene and five 100-gene networks, using Bayesian blocks discretization and the maximum likelihood estimator. The PID-based algorithm PIDC and the raw PUC score perform well in general, as does CLR. All algorithms perform better with larger datasets, but for the larger networks, this improvement is more marked in the algorithms that consider higher order information or network context, suggesting that these are important principles for inferring networks from single cell data. **B)** Dropouts are simulated from the medium-sized 100-node datasets: the lowest 20% (low rate) or 50% (high rate) of expression values for each gene each have a 50% probability of being set to 0. Relative performance of the algorithms is the same in the presence of dropouts, though performance of all algorithms deteriorates with a higher proportion of dropouts. **C)** Violin plots of AUPR scores for all algorithms from all datasets demonstrate the value of higher order information — PUC improves on MI; PIDC improves on CLR — and of network context — CLR improves on MI; PIDC improves on PUC. (x indicates 50-gene network; * indicates 100-gene network; + indicates 100-gene network with dropouts; size indicates number of cells in the dataset). All algorithms used are described in Methods and Results, with MI indicating the use of mutual information scores alone to rank edges (i.e. MI relevance network); the R package minet was used for the existing inference algorithms (Meyer et al. 2008).

##### Box 2: MI-based algorithms

*Relevance networks* (Butte et al. 2000) use the MI estimates (or, in some cases, correlation) in order to detect edges. As there is no reliable universal way of determining the statistical significance of MI values, a threshold is typically chosen to determine which edges are present. This fails to account for the fact that MI may be increased for nodes *X* and *Z* even though they only indirectly interact via an intermediate node *Y*. The *Data Processing Inequality* (DPI) allows us to sort out some of these cases by virtue of the relationship

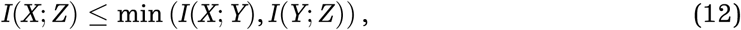

which holds whenever *X*, *Y* and *Z* form a Markov Chain. Post-processing of the MI values using the DPI is at the core of the popular *ARACNE* algorithm (Margolin et al. 2006a, Margolin et al. 2006b). Thresholds on the pairwise MIs are used to identify likely dependent pairs *X*, *Z*; MI values above the threshold are then considered with every possible other node *Y* in light of the DPI.

Given that MI values are affected by a number of factors, including especially the variability of each individual random variable, any global *a priori* threshold may be highly problematic: it will give rise to false positives as well as false-negatives. In the CLR algorithm (Faith et al. 2007, Watkinson et al. 2009) the MI between *X* and *Z* is considered against all MI values for pairings of *X* and *Z* with all other variables *Y*. Thus the threshold for each pair will reflect the variabilities of both genes, as well as their relative levels of statistical dependence on other genes. *MRNET* (Meyer et al. 2007) aims to identify a minimally redundant but maximally explanatory set of variables/predictors for each target gene *X* in a greedy manner.

There have been attempts at using conditional mutual information, interaction information, (4), or related concepts, for network inference (though not applied to single cell data) (Watkinson et al. 2009, Villaverde et al. 2013, Villaverde et al. 2014, Liang & Wang 2008, Zhao et al. 2016). These would have to deal with the known difficulties (Timme et al. 2014) of interpretation (which do not arise in relation to PID-based measures), which may explain the lack of their widespread uptake.

### Application to single-cell data

The extensive analyses using simulated data are necessary to validate our algorithm and provide quantitative comparisons with existing methods. When working with real experimental datasets, we of course do not know the true underlying network, and thus rely on identifying relationships that are consistent with our current biological knowledge about the systems we are studying. Here, we apply our algorithm to three published experimental datasets and, in a related manuscript (Stumpf et al. 2017), illustrate how it can be used as part of a thorough modelling analysis of neuronal differentiation of mouse embryonic stem cells.

Psaila *et al.* (Psaila et al. 2016) used single-cell quantitative PCR (sc-qPCR) to study megakaryocyte-erythroid progenitors (MEP) during human hematopoiesis. Their analysis revealed the existence of subpopulation structure in this class of cells: two groups of cells are primed preferentially for a particular cell fate — megakaryocytic (MK-MEP) or erythroid (E-MEP) — while a third group of multi-potent progenitors (Pre-MEP) retain some myeloid differentiation capacity (Fig. 7A). Here we apply our PIDC algorithm to their complete dataset and infer a candidate network that depicts statistical dependencies among the analysed genes (Fig. 7B). Given we are interested in genes involved in differentiation processes, we also consider networks inferred using overlapping subsets of the data and colour each edge in Fig. 7B according to their presence in these additional networks. Edges that are present in the network constructed using Pre-MEP & E-MEP cells, but not that based on Pre-MEP & MK-MEP cells, are coloured red (i.e. erythroid specific edges), and the reverse (megakaryocytic specific) are shown in blue (edges present in both, or just in the original complete network are shown in grey). Consistent with existing knowledge about these two lineages, we see a cluster dominated by blue edges that comprises known megakaryocytic genes (e.g. *Cd9, Lox, Vwf, Nfib, Cd61, Tgf 1*) and a cluster with several red edges comprising known erythroid genes (e.g. *Cd36, Klf1, Lef1, Cnrip1, Tmod1, Ank1, Dhrs3*) (Psaila et al. 2016). Networks based on pairwise MI scores from the same data show skewed degree distributions with many nodes unconnected (Fig. 7C-D).

**Figure 6:**
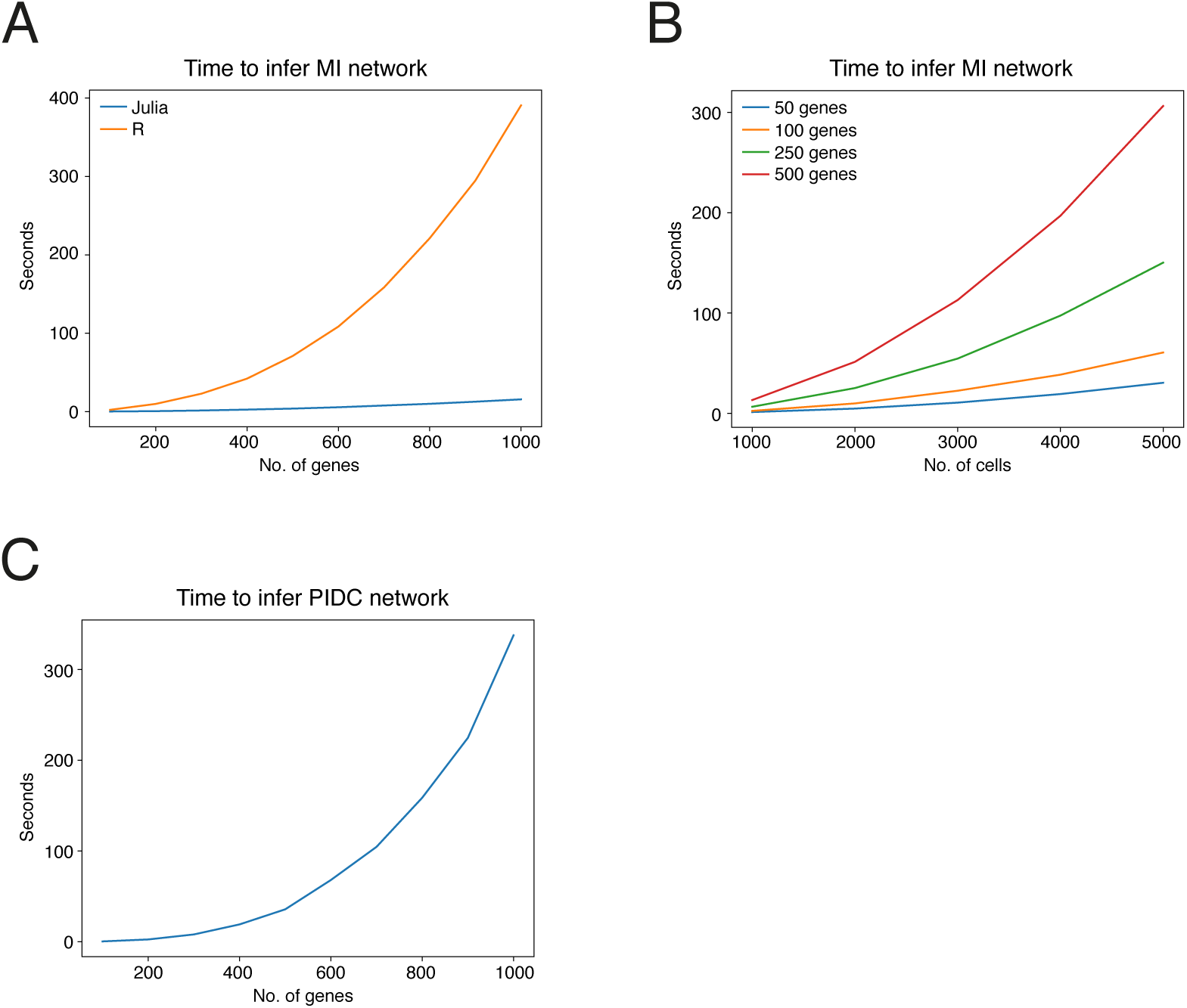
Speed of MI and PIDC calculations using the Julia programming language. **A)** Comparison of the times taken to calculate a matrix of pairwise MI values using the R package minet (Meyer et al. 2008) and our Julia package InformationMeasures.jl. Input data were simulated expression values for up to 1000 genes, with 700 values per gene (equivalent to 700 cells, the same as our medium-sized in silico dataset). Data were discretized using the uniform width algorithm, because Bayesian blocks was not available via minet, and times were measured using inbuilt functions in R and Julia. **B)** Times taken to calculate the MI matrix in the Julia programming language with larger numbers of cells, for networks of different sizes. Data were discretized using the recommended Bayesian blocks algorithm, which has a much greater complexity than the uniform width algorithm, but produces better estimates. **C)** Time taken to infer networks of varying sizes using the PIDC algorithm implemented in Julia. Networks were inferred for simulated datasets of up to 1000 genes, with 700 expression values per gene.

**Figure 7:**
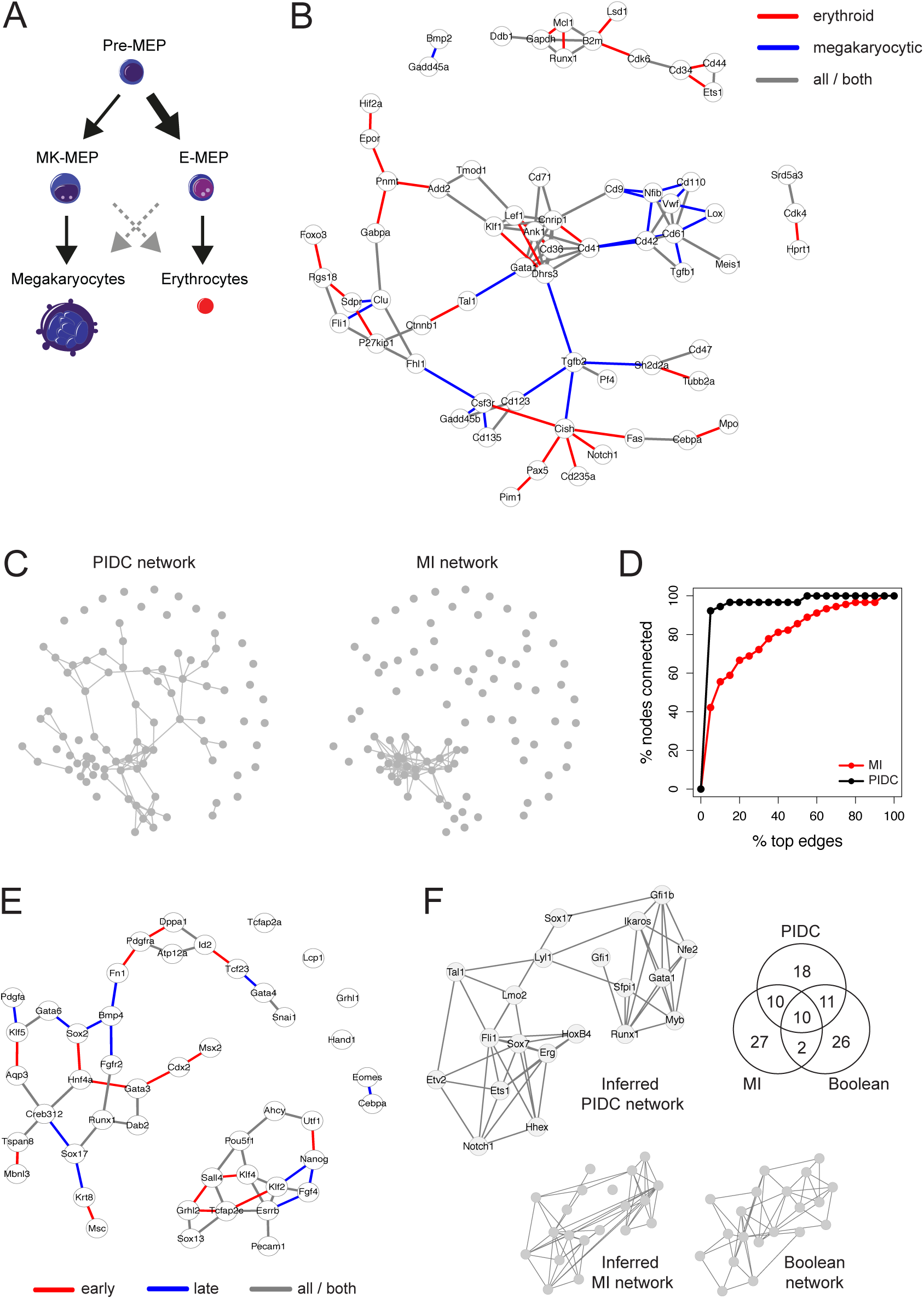
Application of the PIDC inference algorithm to experimental datasets. **A)** Illustration of the relationship between the three subpopulations of MEP cells — ‘Pre-MEP’ cells are enriched for erythroid/megakaryocyte progenitors but still retain some potential to differentiate into other cell types (myeloid cells); ‘E-MEP’ and ‘MK-MEP’ cells are strongly biased towards erythroid and megakaryocyte differentiation respectively (for details see Psaila et al. (2016). **B)** Network inferred using the PIDC algorithm from the complete set of data from Psaila et al. (the top 2.5 % of edges are shown; for clarity, only nodes connected by these edges are shown). Edge colours indicate whether these edges are also detected in networks constructed using subsets of the data (comprising data from Pre-MEP cells combined with either E-MEP or MK-MEP cells). Red edges indicate those that are present in the Pre-MEP & E-MEP network but not the Pre-MEP & MKMEP network (i.e. erythroid specific), while blue edges indicate the reverse scenario (i.e. megakaryocytic specific); edges present in both networks, or only in the network constructed using all the data, are shown in grey. **C)** Comparison of the networks inferred from the data from Psaila et al. using our PIDC algorithm or MI relevance networks (in both cases, the top 2.5 % of edges are shown). **D)** Percentage of nodes that are connected in networks inferred from the Psaila et al. data as the threshold for edge inclusion is varied (from 0 to 100 % of possible edges, according to their rank). These results show that networks inferred using PIDC (black) tend to be better connected than networks inferred using MI (red); i.e. MI networks show more skewed degree distributions. **E)** PIDC interaction network inferred using early embryonic development data (oocyte to E4.25 blastocyst stages), see Guo et al‥ Graph edges indicate the top 5 % of putative interactions detected using the PIDC algorithm on the complete dataset. Networks are also inferred using two overlapping subsets of the data — an ‘early’ subset that includes all cells collected from oocyte up to 32-cell E3.5 blastocyst stages, and a ‘late’ subset including cells collected from 16-cell morula to 64-cell E4.25 blastocyst stages. Edge colours indicate temporal dependencies of the identified relationships; red indicates an edge ranks in the top 5 % of edges in the early network but not the late network, blue indicates the converse (late but not early), while grey indicates relationships without specific temporal dependencies (i.e. only present in the network constructed using the complete dataset, or present in both the early/late networks). **F)** Comparison of networks inferred using haematopoietic development data in Moignard et al‥ The authors used single-cell expression data for 20 transcription factors to infer a Boolean network model of blood development; we show a simplified representation of their model, where nodes (genes) are linked by an edge if those genes are either directly linked, or linked via one Boolean operation or set of update rules (genes linked in their model via a chain of multiple sets of update rules are not connected here). We used our PIDC algorithm or MI alone to infer networks of putative interactions between genes using these same data, and compare the edges identified in each of these three networks (numbers of shared edges are indicated by the Venn diagram).

We next consider a sc-qPCR dataset comprising expression measurements of selected genes during early embryonic development (from oocyte to 64-cell blastocyst stages) (Guo et al. 2010); Fig. 7E shows the resulting inferred network. We again infer additional networks using subsets of these data — here we use overlapping subsets of ‘early’ and ‘late’ cells to reveal any temporal dependencies in the detected interactions. A number of known relationships between genes are apparent in the network, e.g. upregulation of *Cdx2* and *Gata3* transcription factors (TFs) during the 8-cell to morula transition is identified as an edge in the ‘early’ network; while the co-expression of primitive endoderm specific TFs *Creb312* and *Sox17* is detected as an edge in the ‘late’ network (consis-tent with the appearance of distinct cell types, including primitive endoderm cells, in the blasto-cyst). A cluster of known pluripotency and reprogramming factors is also identified in the network (*Pou5f1, Nanog, Esrrb, Klf2* and *Klf4*) — *Sox2*, another key reprogramming factor, is not connected with these genes but is known to be up-regulated later than the other factors (Guo et al. 2010).

Moignard *et al.* studied embryonic haematopoietic development, and used sc-qPCR data to develop a Boolean network model of the GRN underlying blood development (Moignard et al. 2015). In Fig. 7F we compare the networks inferred from these same data using PIDC and MI — we find the inferred PIDC network shares a higher number of edges with the Boolean network model, than the network constructed using MI values alone. Although we of course do not know the ‘true’ structure of the GRN in this case (this is only feasible when using *in silico* data), the Boolean model was shown to capture key cell states observed experimentally and generated several experimentally-validated predictions (Moignard et al. 2015), thus we use this as a benchmark to indicate the biological plausibility of our inferred networks.

In a related companion paper (Stumpf et al. 2017) we apply our PIDC algorithm to sc-qPCR data collected from cells undergoing differentiation from a pluripotent ground state towards a committed neuronal lineage, via a primed epiblast-like state. 547 cells were sampled in total at seven time-points spanning seven days; we analyse expression measurements of 74 genes including known regulators of pluripotency and neuronal differentiation (see (Stumpf et al. 2017) for details). We first assign the cells to three robust groups that correspond with developmental stage (using k-means clustering), and then infer networks using data from all the cells, or from overlapping subsets containing cells at earlier or later stages of development. Comparing the networks obtained using different subsets of cells allows us to observe any temporal dependencies in the inferred interactions. Using an unsupervised community detection algorithm, we find that the network of inferred (co-)regulatory relationships contains several communities (or modules) of genes displaying high connectivity within each community. Three of these communities show distinct temporal dependencies in connectivity, and comprise genes known to play roles at different stages of differentiation. Our analysis thus identifies modules of genes that undergo coordinated changes in expression as cells progress through development, and putative gene interactions that may be involved in regulating these transitions in cell state.

### Guidelines & limitations

Any comparative analysis of information-based GRN inference algorithms is influenced by a number of decisions, in particular: (i) how the data are discretized, (ii) the choice of MI estimator, and (iii) the metric used to evaluate performance. We discuss the impact of each of these decisions and offer guidelines for future analyses, before discussing the use of single cell RNA sequencing (scRNA-seq) datasets.

The information measures described here all rely on estimates of discrete probability distributions. Normalised mRNA expression data are generally continuous, but estimating the distributions for continuous random variables is fraught with problems. Several algorithms and heuristics have been developed to discretize data and estimate empirical probability distributions from the resulting discrete frequencies. We investigated two methods for discretization, along with four MI estimators, as described in *Methods*.

All estimators produce fairly accurate estimates of joint entropies for up to two uniformly distributed random variables, but in higher dimensions performance varies according to the distribution (Tables S2 and S3), making it difficult to identify the most appropriate estimator for experimental data. Rank agreement between the estimators is good when the data are discretized using Bayesian blocks, however, diminishing the importance of the choice of estimator (Fig. S4). In light of these findings we advise using Bayesian blocks (an adaptive discretization algorithm that allows variable width bins) and since the underlying distribution is usually unknown we favour the maximum likelihood estimator due to its simplicity.

The choice of discretization method and estimator influences the performance of the inference algorithms, with effects varying depending on the algorithm and on the true network (Fig. S3). Sampling frequency and dataset size also have an effect (Fig. 5A), with performance increasing in line with data set size, and decreasing in larger networks and with a higher rate of technical dropout errors (Fig. 5B). Due to the number of influential factors, we advise caution when interpreting the results of this or any such comparison as an exhaustive exploration of these factors is not feasible; however we note that the PIDC algorithm performs well in general across the many tested combinations of discretization methods, estimators and datasets.

The metric used to evaluate algorithms also affects their apparent performance, evident here in the higher scores for AUROC than AUPR (Fig. S3B). This is a well-documented phenomenon, caused by the true negatives (unconnected node pairs) in a GRN vastly outnumbering the true positives (edges); for example, the *E. coli 1* 100-gene network contains 125 edges and 4825 unconnected pairs (Table S1). AUROC equally rewards the prediction of an edge and a non-edge, meaning that the score for any algorithm that mostly (or even exclusively) predicts non-edges will be misleadingly inflated, however well or badly it predicts edges. AUPR is therefore the more meaningful measure, despite AUROC being widely-used (Murphy 2012).

Here, we have illustrated the application of our method using several single cell qPCR datasets; scRNA-seq experiments generate much larger datasets comprising expression measurements for thousands of genes. When analysing these data, a subset of (up to hundreds of) genes should first be selected — both to make the network inference analysis computationally tractable (see *Software*), but also to aid interpreting the results. There are many potential approaches to selecting gene subsets, depending on the purpose of the analysis and the level of existing knowledge about the specific system being studied. We may wish to select genes likely to be involved in the process of interest based on prior knowledge (and/or functional annotations of genes) such as known transcription factors, similar to the way that genes are selected for analysis in qPCR experiments. However, we can also make use of the huge array of statistical and computational methods that have been developed to analyse scRNA-seq experiments and select subsets of genes based on their observed gene expression patterns (Bacher & Kendziorski 2016, Stegle et al. 2015, Grün & van Oudenaarden 2015, Liu & Trapnell 2016). For example, we could select those showing higher than expected levels of expression variability, those that show differential expression between cell states or over time (using the results of clustering or pseudotemporal ordering algorithms), or cluster genes by similar expression profiles and select representative genes from each cluster. As is the case for other network inference algorithms, genes with no variability in mRNA expression are uninformative, and should always be removed prior to analysis. In addition, due to the prevalence of zeros and lack of sensitivity of single cell experiments, many genes (particularly those expressed at low levels) will not be reliably detected so we can also exclude those without detectable expression in a large proportion of cells.

### Software

A new open-source package for estimating MVI measures is implemented in the Julia programming language (Bezanson et al. 2014). The package, named InformationMeasures.jl supports information measures such as entropy, MI, CMI, and PID; the maximum likelihood, Miller-Madow, Dirichlet and shrinkage estimators; and the Bayesian blocks, uniform width and uniform count discretization methods.

Julia was chosen for its speed (Fig. 6A), clear mathematical syntax, growing availability of libraries, and good integration with other languages. The existing Discretizers.jl package is used to implement the discretization methods; in some of our initial analyses we used the AstroML Python implementation of the Bayesian blocks algorithm (Vanderplas et al. 2012, Scargle et al. 2013). In order to meet a wide range of requirements, the package can be used simply for discretizing data, or to calculate information measures using pre-discretized data or probability distributions that have been estimated elsewhere.

A Julia package implementing PIDC and other inference algorithms is available, along with tutorials and our simulated datasets at https://github.com/Tchanders/network_inference_tutorials. Our algorithm has complexity *O*(*n*^3^) in the number of genes, but the speed of our Julia implementation means that inference time is comparable to widely-used implementations of the lower-complexity algorithms (Fig. 6A). The complexity in the number of cells depends on the discretization method: the recommended Bayesian blocks method scales less well than the uniform width method, but nevertheless produces results for several thousands of cells for a network of hundreds of genes within a practically useful timescale (Fig. 6B-C).

## Discussion

Here, we have introduced a network inference algorithm based on PID (Williams & Beer 2010, Timme et al. 2014), an easily interpretable MVI measure that allows us to explore statistical dependencies between multiple genes in detail. Our PIDC algorithm identifies putative functional relationships between genes based on the unique contribution to pairwise MI (Eq. 5) combined with information about the local network context of each gene. We use extensive performance comparisons (Fig. 5) to demonstrate the value of using both higher order information measures and network context, and to illustrate that the large sample sizes provided by single cell data are critical to our algorithm’s success. Like other studies (Simoes & Emmert-Streib 2011, Olsen et al. 2009, Hausser et al. 2009), we find that the methods chosen to discretize data and estimate entropies and probability distributions affect algorithm performance considerably (Fig. S3) — too often, the impact of these choices has been ignored. A fast, open-source software package provides an easy way for users to explore such factors when applying our method.

Although single cell data have many potential advantages over bulk transcriptomic data for network reconstruction approaches — particularly sample size, inherent variability, and ability to detect subpopulation structure — they also pose notable challenges. Technical noise and other biological sources of heterogeneity (e.g. transcription stochasticity, other cellular processes) can, to some extent, impede our ability to detect informative statistical dependencies; there are clear theoretical benefits of using information theoretical methods when we expect the observed dependencies to be complex and non-linear. The relative contributions of different sources of noise and variation are still poorly understood and good noise models are lacking (Bacher & Kendziorski 2016, Fu et al. 2016). We therefore relied on a simple model of ‘dropout’ events to mimic the zero-rich nature of these data in our simulations — as expected, technical noise reduces our ability to recover the true network structure (Fig. 5B). The best performing algorithms, PIDC and CLR, both aim to capture the most important interactions for each node in turn, rather than the highest dependencies across the whole dataset. This leads to well connected inferred graphs (e.g. Fig. 7C), but can also help to address the influence of potential confounding factors (such as the cell cycle) when working with single-cell data. Variation in cell-cycle stage causes large-scale changes in cell transcriptional states (Buettner et al. 2015, Scialdone et al. 2015, McDavid et al. 2014) — and we would expect it to induce stronger statistical dependencies amongst the affected genes. However, using empirical distributions to take network context into account (as in PIDC and CLR) will at least partly mitigate the influence of any such confounding factors.

Integrating our method with other single cell analyses allows us to select subsets of cells and genes that are most informative about our specific biological questions. Firstly, there are many sources of biological heterogeneity in single-cell data so it is important to focus on the variation of interest, e.g. when studying developmental processes we should aim to analyse collections of data where we expect cell differentiation to be the major source of variation. We can use established methods for analysing the subpopulation structure of single cell data — such as clustering, dimensionality reduction, and pseudotemporal ordering algorithms (Bacher & Kendziorski 2016, Stegle et al. 2015, Grün & van Oudenaarden 2015, Liu & Trapnell 2016) — e.g. to identify distinct cell subtypes or alternative differentiation pathways. Here we have shown how this allows us to focus on functional relationships involved in developmental transitions, and the relative timing of transcriptional changes. For example, in (Stumpf et al. 2017) we use clusters of developmentally similar cells to examine how the activity of (co-)regulatory relationships changes during development, making initial steps towards defining mechanistic models of neural differentiation — the network structure and temporal information allow us to suggest candidate genes for maintaining cell states or driving developmental transitions. Although scRNA-seq generates data for thousands of genes, we recommend selecting meaningful subsets for network inference — either based on biological knowledge, or by using gene expression variability and patterns (Bacher & Kendziorski 2016, Trapnell et al. 2014, Haghverdi et al. 2016, Reid & Wernisch 2016, Setty et al. 2016, Kharchenko et al. 2014, Korthauer et al. 2016, Vallejos 2016, Finak et al. 2015). Data imputation methods, e.g. (van Dijk et al. 2017), or average expression measurements over small groups of similar cells may help addressing the challenges of noise and low coverage.

For any statistical approach it is important to consider the potential limitations. Many are general to network inference approaches aiming to reconstruct GRN structure from mRNA profiles and are discussed in depth elsewhere (Penfold & Wild 2011, Marbach et al. 2012, Villaverde & Banga 2013, Oates & Mukherjee 2012), but it is worth emphasising a few key points that affect how we should interpret and use our results. Firstly, we can only detect relationships where there is sufficient variability in gene expression observed under the chosen experimental conditions. Functional interactions are only detectable if they induce changes in transcriptional state that persist over a reasonable timescale — we will not, for example, detect rapidly fluctuating changes, as the transient changes in mRNA levels will not result in observed statistical dependencies across cells. As well as functional regulatory relationships, we are likely to also identify co-regulatory relationships where genes under the influence of the same regulator show coordinated expression changes. Without making further assumptions, or using perturbation or temporal data, we cannot distinguish *causal* relationships; however, in many instances it will still be informative to learn which sets of genes respond in a coordinated manner. In cases where the assumptions made by pseudotemporal ordering algorithms (Trapnell et al. 2014, Bendall et al. 2014, Haghverdi et al. 2016, Reid & Wernisch 2016, Setty et al. 2016, Moris et al. 2016) are justified, we can potentially use this information to infer causality and directionality of gene interactions (Villaverde et al. 2014, Villaverde et al. 2013, Zoppoli et al. 2010, Opgen-Rhein et al. 2007). It is of course unrealistic to expect every cell to follow precisely the same route through transcriptional space; we do instead make the modest assumption, that there are key changes in transcriptional state that must occur in order for cells to respond appropriately to environmental and developmental cues, and that these will be subject to conserved regulatory mechanisms.

Methods for exploring high-throughput single cell datasets and identifying putative functional relationships between genes are clearly needed. As with all network inference methods, we cannot expect to reconstruct the exact structure of the underlying biological networks, but instead view such methods as tools to explore the data; generate hypotheses; represent the current state of understanding; and guide further experiments, model development and analyses. Validating or invalidating these hypotheses experimentally may, of course, lead to revised network models — like any mathematical model it should be subject to refinement as new facts are assembled and new insights are gained.

## Methods

### Discretization algorithms

In order to use the entropy estimators described here, continuous datasets must first be discretized. A number of algorithms exist to define the total number and boundaries of the resulting partitions (or bins). One common simple approach is to use bins of equal width, with the number of bins determined heuristically, e.g. here we use the nearest integer to the square root of the size of the dataset, *p*_*n*_ (Mosteller et al. 1977). A more sophisticated approach is the Bayesian blocks algorithm (Scargle et al. 2013), in which the number and widths of bins are chosen by optimising a fitness function, without constraining the bins to be of equal width.

### Estimators

When a dataset is large enough, the empirical frequencies can be considered to be an approximation of the true probabilities, referred to here as the maximum likelihood approach. For sparser datasets, a number of methods have been developed either for estimating the probability distribution from a set of frequencies, such as the Dirichlet estimator and the shrinkage estimator, or for estimating the entropy directly, as in the Miller-Madow estimator (Hausser & Strimmer 2009, Paninski 2003).

The Dirichlet estimator refers to a group of Bayesian estimators that take a Dirichlet distribution as prior, but each with different parameters (Agresti et al. 2005, Hausser & Strimmer 2009). There is no consensus on the best parameters to use, despite several proposed alternatives (Hausser & Strimmer 2009); here, we use the same parameter, 1, for each bin unless otherwise stated.

The shrinkage estimator is also Bayesian, compromising between the observed frequencies, unbiased but with a high variance, and a prior (or target) distribution, biased but with low variance (Hausser & Strimmer 2009). The estimate is affected by both the choice of target distribution and the weight given to the target (or shrinkage intensity). In the current analysis the optimal shrinkage intensity is calculated as described in (Hausser & Strimmer 2009), and the target distribution is the uniform distribution.

The Miller-Madow estimator is an entropy bias correction that does not estimate the probability distribution, and therefore cannot be meaningfully applied to higher order information measures. Despite this it has been applied for the comparison of different MI-based algorithms (Meyer et al. 2008), and so it is included in this analysis, with the caveat that its meaning is unclear.

### Simulation of *in silico* 3-gene network data

We considered six 3-gene topologies (Fig. S1), and used the Gillespie algorithm (Gillespie 1977) to generate stochastic simulations of gene expression timecourse data using two alternative model definitions (based on thermodynamic or mass action kinetics). In both cases, we include an additional activating stimulus that is present from halfway through the simulation timecourse. This additional stimulus acts to perturb the system away from a steady state, driving changes in gene expression that are necessary for relationships between genes to be observable. This section describes these model and simulation details.

The first *thermodynamic* model includes seven species: mRNA (*x*_*i*_) and protein (*y*_*i*_) corresponding to three genes (*i* = 1, 2, 3), and a stimulating ligand (*s*) which targets a selected gene. We define the following reaction types with associated propensities,

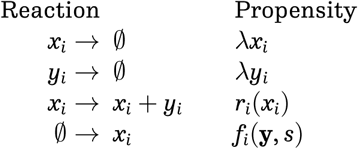

to represent mRNA decay, protein decay, translation and transcription respectively, where is the protein / mRNA decay rate, translation is modelled according to saturation kinetics (with maximum rate α _translation_), i.e.,

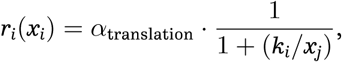

and transcription rates depend on the concentration of any regulating proteins (including the stimulating ligand *s* if present) according to the relationship,

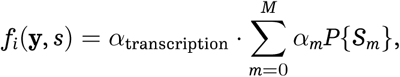

where *α*_transcription_ is a constant transcription rate, *M* is the total number of possible states *S*_*m*_ for gene *i* (either unbound, or bound by one or two regulating proteins), and *α*_*m*_ is the relative activation rate for each state. The probability of each state, *P*{*S*_*m*_}, depends on the concentrations of the regulating proteins, modelled according to standard thermodynamic principles (see e.g. (Marbach et al. 2010) for details). For example, if a gene has two possible regulators (proteins *y*_*j*_, *y*_*k*_), we calculate the mean activation of transcription of the target gene *i* using the function,

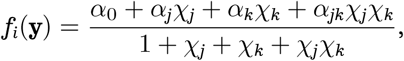

where *χ*_*j*_ = (*y*_*j*_/*k*_*j*_), *k*_*j*_ is the dissociation constant, and the possible states of the gene are unbound, bound by *y*_*j*_ or *y*_*k*_ alone, or by both *y*_*j*_ and *y*_*k*_. For our models, the maximum number of regulators for a given gene is three (proteins *y*_*j*_ and *y*_*k*_ plus the stimulating ligand *s*), but we assume a maximum of two regulators can bind the gene at any one time (so we consider the gene states with each possible pair of ligands bound, but not a state with all three bound).

For the thermodynamic model, we simulated timecourses from time 0 to 1000, with the stimulating ligand *s* present from time 500 at a constant level of 20 molecules, and recorded the system state at 41 equally spaced intervals. We repeated the simulations 25 times — and calculated PID scores using the resulting data (i.e. 1025 data points or ‘cells’ were used to calculate each PID measure)— with the stimulus targeting each of the three genes in the network in turn (75 simulations in total, 3 sets of PID scores). We randomly sample initial mRNA (*x*_*i*_) and protein (*y*_*i*_) levels from a *U*(0, 5) distribution; we perform simulations for five different initial conditions and plot the mean PID profiles from these five different conditions in Figs. 2A and S1. Model parameters are = 0.02, α_transcription_ = 2, α_transcription_ = 2, and *k*_*i*_ = 50 (for all *i* = 1, 2, 3). Relative activation rates for transcription, α_*m*_, depended on the number of activating and inhibiting regulators present in each possible state *S*_*m*_: α_*m*_ = 0.1 (for the unbound state, i.e. basal transcription), 0.001 if an inhibitor was bound (we assumed inhibition dominated activation), and 5 if only activating regulators are bound.

The second *mass action* model that we consider also includes seven species: genes (*g*_*i*_) and mRNA/protein (*x*_*i*_) for *i* = 1, 2, 3, and the stimulating ligand *s*. We assume that protein and mRNA concentrations are equal (i.e. translation is instantaneous) and, unlike our first model, assume that a gene can only be bound by a single protein at any time. The possible reactions and associated propensities are,

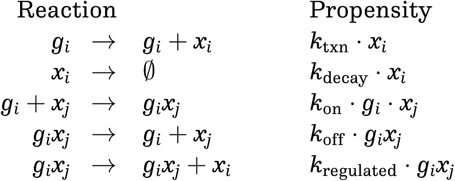

for basal transcription and protein decay, protein binding and unbinding from a target gene, and transcription from a gene bound to a regulating protein respectively (where *g*_*i*_*x*_*j*_ indicates gene *i* is bound by regulating protein *j*).

For the mass action model, we simulated timecourses from time 0 to 400, with the stimulating ligand present from time 200 at a constant level of 20 molecules, and recorded the system state at 21 equally spaced timepoints. We repeated the simulations 50 times with the stimulating ligand targeting each gene to give a total of 1050 data points (‘cells’) that are used to calculate PID measures. We initiated simulations with two copies of each gene (*g*_*i*_ = 2), no stimulating ligand, and initial mRNA/protein levels sampled from a uniform distribution (*x*_*i*_ ~ *U*(0, 50)); we perform simulations for five different initial conditions and calculate the mean PID profiles from these five conditions (plotted in Figs. 2A and S1). Model parameters are *k*_txn_ = 1, *k*_decay_ = 0.05, *k*_on_ = 0.01, *k*_off_ = 0.25, and *k*_regulated_ = 10 or 0.1 for activating and inhibiting regulation respectively. Information measures were calculated from these data using the Matlab package written by Timme *et al.* (Timme et al. 2014), following discretization with the AstroML implementation of the Bayesian blocks algorithm (Vanderplas et al. 2012, Scargle et al. 2013).

### Simulation of *in silico* GeneNetWeaver network data

Data are simulated using GeneNetWeaver (Schaffter et al. 2011), a software package which generates stochastic simulations from *in silico* networks that are designed to be representative of real biological network structures (they are created by extracting subnetworks from known *E. coli* and *S. cerevisiae* transcriptional networks). This software has become a common tool for simulating gene expression data — including its use as part of several DREAM (Dialogue on Reverse Engineering Assessment and Methods) network inference competitions (Schaffter et al. 2011, Marbach et al. 2010, Marbach et al. 2012) — which aims to provide unbiased datasets that do not favour particular inference methods, and networks that retain characteristics of real GRNs. We compare the network inference algorithms using ten networks, five with 50 genes and five with 100 genes; for each network size there are two *E.coli* and three *S.cerevisiae* networks, with average node degrees ranging from 1.19 edges per node to 5.51 edges per node.

GeneNetWeaver uses dynamical models that consider mRNA transcription and translation processes and generates time series simulations using stochastic differential equations to model dynamical noise and a mixed normal and log-normal model to represent microarray noise. To mimic single-cell data, we simulated thousands of time series experiments for each network, using the default settings, with mRNA measurements generated according to the default settings and the default time points: times 0 to 1000, in steps of 50. We sampled a single time point from each time series, representing a single ‘cell’: for the large datasets, we sampled 100 cells from each of the time points (2100 cells in total); for the medium datasets we sampled 100 cells from time 0 onwards in time steps of 150 (700 cells); and for the small datasets, we sampled 20 cells from all time points between 0 and 300 (140 cells). Where the dataset size is unspecified, we have used the medium datasets. For our initial exploration of PID measures, we also used data simulated from a 10-gene network (shown in Fig. 3A); in this case we again used the temporal sampling scheme for medium datasets.

In order to test the robustness of our method to zero-inflated data, typical of single-cell experiments, we further simulated ‘drop-out’ datasets. Zero measurements appear to be a combination of technical errors and genuine lack of expression due to stochasticity or biological state, and are more common in transcripts with low abundance (Kharchenko et al. 2014, Brennecke et al. 2013). We simulated drop-out events at two rates, such that expression values in the lowest 50 % (high rate) or 20 % (low rate) for each gene had a 50 % probability of being recorded as 0.

### Network inference performance metrics and comparisons

AUROC and AUPR curves are calculated by comparing the inferred networks (which assign a score to every potential network edge) to the ‘true’ network used to simulate data, and identifying the numbers of correctly (and incorrectly) assigned edges as the threshold for edge inclusion is varied. AUROC is calculated from the area under the ROC curve, which is a plot of the false positive rate (FPR) on the *x*-axis versus the true positive rate (TPR) on the *y*-axis. AUPR is the area under the curve for a plot of precision (*y*-axis) versus recall (equal to TPR) on the *x*-axis. These quantities are calculated as,

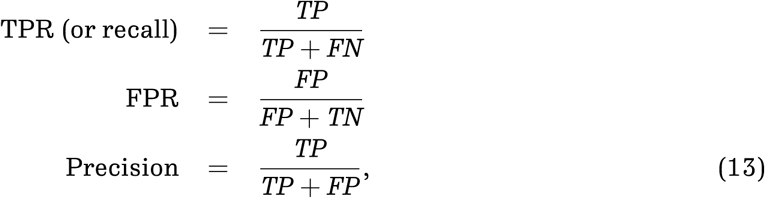

where *TP* and *FP* indicate the numbers of true and false positives, and *TN* and *FN* are true and false negatives. For networks where the number of negatives is much greater than the number of positives, AUPR is considered a better metric for comparing algorithm performance (Murphy 2012, Davis et al. 2006).

We use these scores to compare performance of our method relative to existing inference algorithms (Figs. 5 and S3). We used the implementations in R package minet for the existing inference algorithms (Meyer et al. 2008); we used the default or suggested parameters within this package except for the tolerance, *τ*, for ARACNE for which we either used the default (for the results in Fig. S3) or 0.1 (for the results in Fig. 5, as recommended in the original publication describing this algorithm (Margolin et al. 2006a)).

## Methods for analysis of real datasets

### Published datasets

We analysed three published qRT-PCR datasets to illustrate our network inference algorithm. Normalised Ct values from Psaila *et al.* (Psaila et al. 2016) were subtracted from the assumed maximum, 40, and the resulting dCt values (for 87 genes and 681 cells) were used in our analyses. Normalised dCt values from Moignard *et al.* (Moignard et al. 2015) are used directly for our analyses; we used data from the 20 genes they represent in their network model and 3934 cells. Raw Ct data from Guo *et al.* (Guo et al. 2010) are treated as described by the original authors (dCt values are calculated assuming a limit of detection of 28, and normalised on a cell-wise basis by subtracting the mean expression of housekeeping genes *Actb* and *Gapdh*; all values corresponding to expression below the limit of detection are set to −15); we used data from 46 genes (i.e. we excluded the housekeeping genes used for normalisation) and 442 cells.

## Availability of data and materials

The InformationMeasures.jl package is available from https://github.com/Tchanders/InformationMeasures.jl.

A Julia package for running the PIDC, PUC, CLR and MI algorithms is available from https://github.com/Tchanders/NetworkInference.jl.

Tutorials and simulated datasets are available from https://github.com/Tchanders/network_inference_tutorials.

## Author’s contributions

TEC, MPHS, and ACB designed and performed research, analysed data, and wrote the paper. All authors read and approved the final manuscript.

## Acknowledgements

This work was supported by a Biotechnology and Biological Sciences Research Council (BBSRC) DTP Studentship to TEC, and a BBSRC Future Leaders Fellowship to ACB. We thank Joe Greener, Gal Horesh, and Ananth Pallaseni for sharing code with us, Suhail Islam for computing support, and Ben MacArthur and Patrick Stumpf, as well as the members of the theoretical systems biology group for useful discussions.

## Supplementary Information for Gene regulatory network inference from single-cell data using multivariate information measures

**Figure S1:**
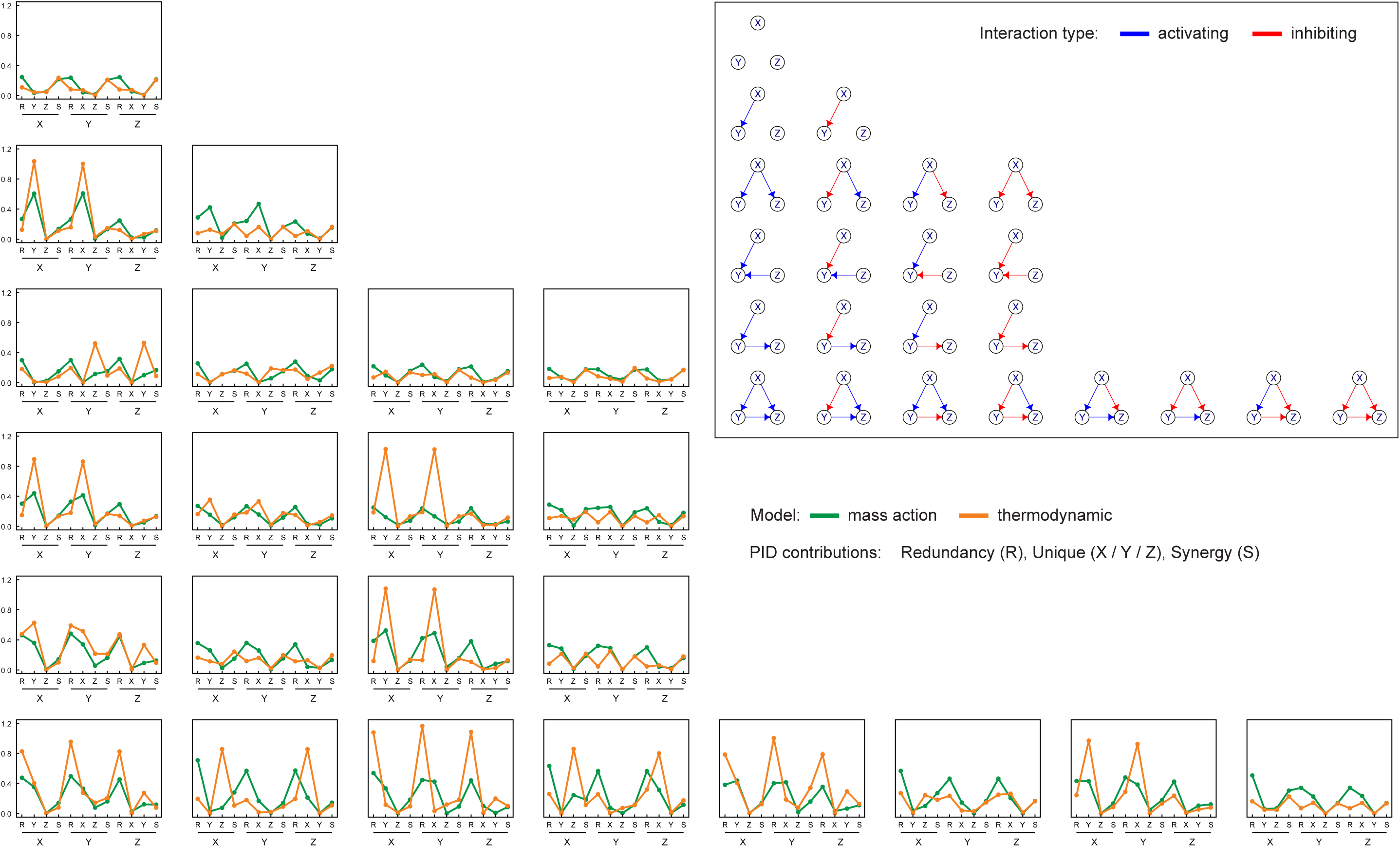
Related to Figure 2. Mean PID profiles for 3-gene network simulations, with stimulating ligand targeting gene X, and different network topologies. PID values were calculated using data simulated from 3-gene networks with the topologies illustrated in the top right inset (blue and red arrows indicate activating and inhibiting interactions respectively). Each line graph in the main figure shows the mean PID values calculated using models with the topology indicated in the equivalent grid position (i.e. the same row/column); the horizontal axis labels indicate the PID contribution, e.g. the first four values show the PID values with gene X as the target, consisting of the redundancy (R), unique contributions from gene Y (Y) and gene Z (Z), and the synergistic contribution (S). The models used for simulation assumed mass action (green) or thermodynamic (orange) kinetics, and the additional stimulation ligand targeted gene X from halfway through the simulation time (see Methods). The values plotted are the mean PID values calculated from five sets of simulations (with different randomly sampled initial conditions). Note that the same profiles are not necessarily seen for networks with equivalent connectivity, but different types of edges (results in the same row of this figure) | this is to be expected, as statistical relationships between genes can only be detected when there is sufficient variability in the observed data (which may not occur in some cases, e.g. if expression of a gene is inhibited throughout the simulation time).

**Figure S2:**
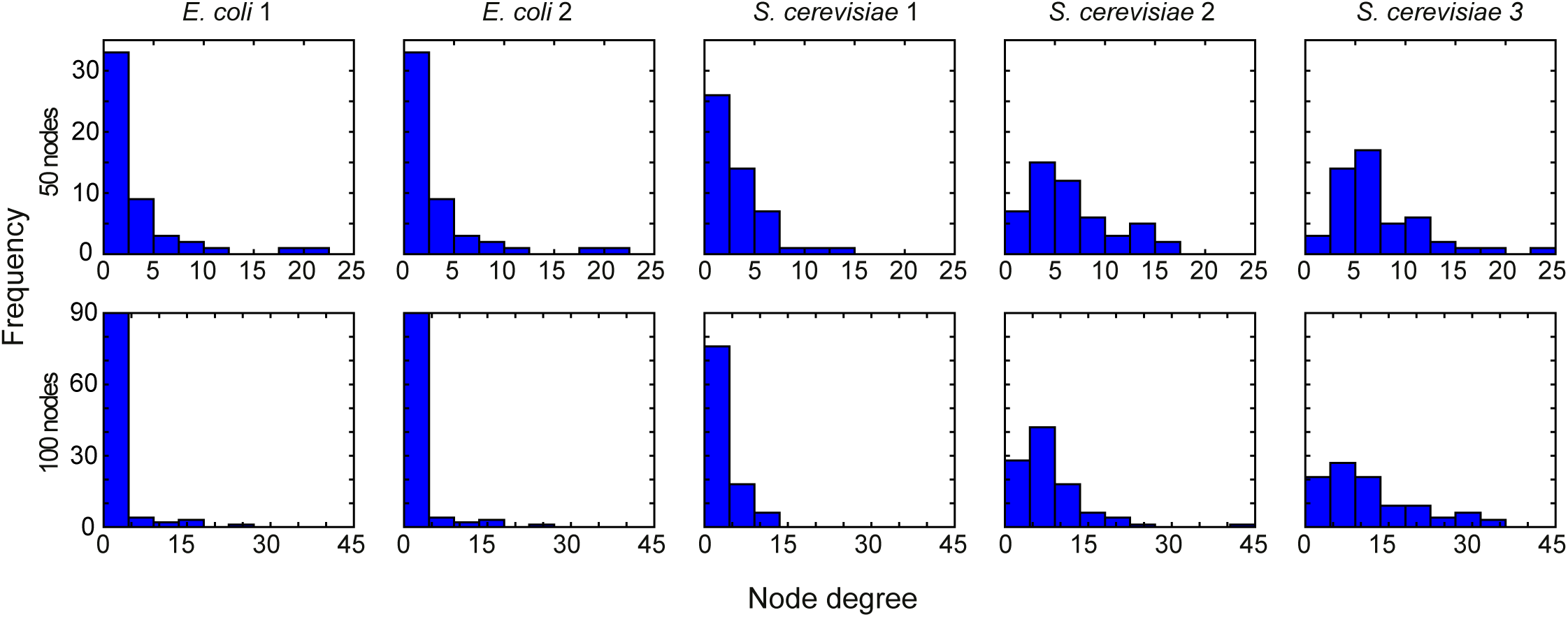
Related to Figure 2 and Table S1. Node degree distributions for each of the 50-node and 100-node networks produced by GeneNetWeaver (Schaffter et al. 2011). The distributions are varied, with the *E. coli* networks tending to have more hubs, and the *S. cerevisiae* networks tending to be more densely connected.

**Figure S3:**
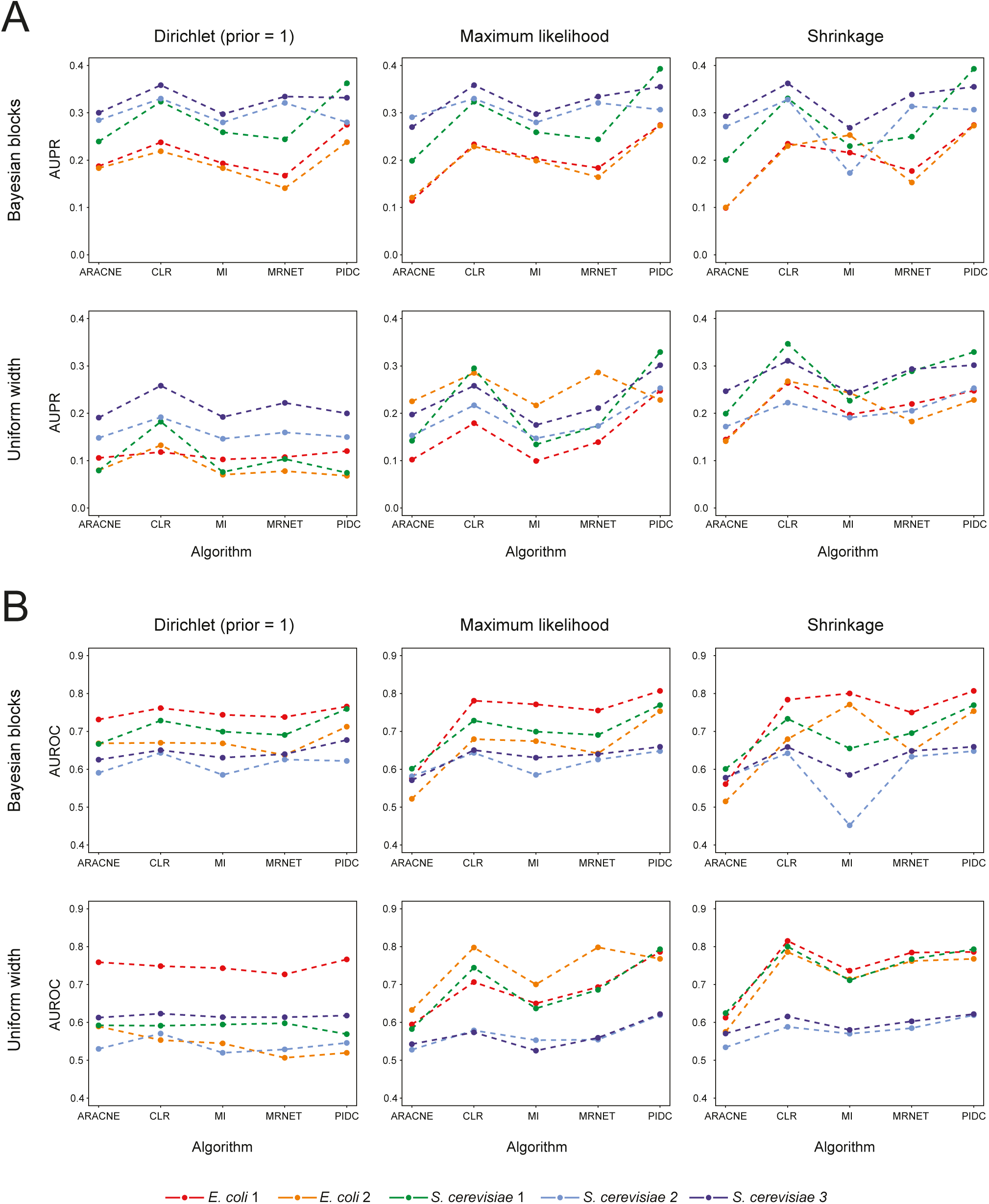
Related to Figure 5. Influence of discretization algorithm and estimator on network inference performance. AUPR (A) and AUROC (B) scores quantifying the accuracy of inferred networks calculated using *in silico* data simulated from five 50-gene networks (see *Methods*). Coloured lines indicate the results obtained with different datasets. Each plot shows the results obtained using a different combination of discretization algorithm (rows) and MI estimator (columns). Choice of algorithm and estimator clearly affects the relative scores of the network inference algorithms, suggesting that comparisons made using inconsistent combinations should be interpreted with caution; we find that the PIDC inference algorithm performs consistently well across the different combinations. The R package minet was used to implement the existing algorithms using the default parameters (Meyer et al. 2008).

**Figure S4:**
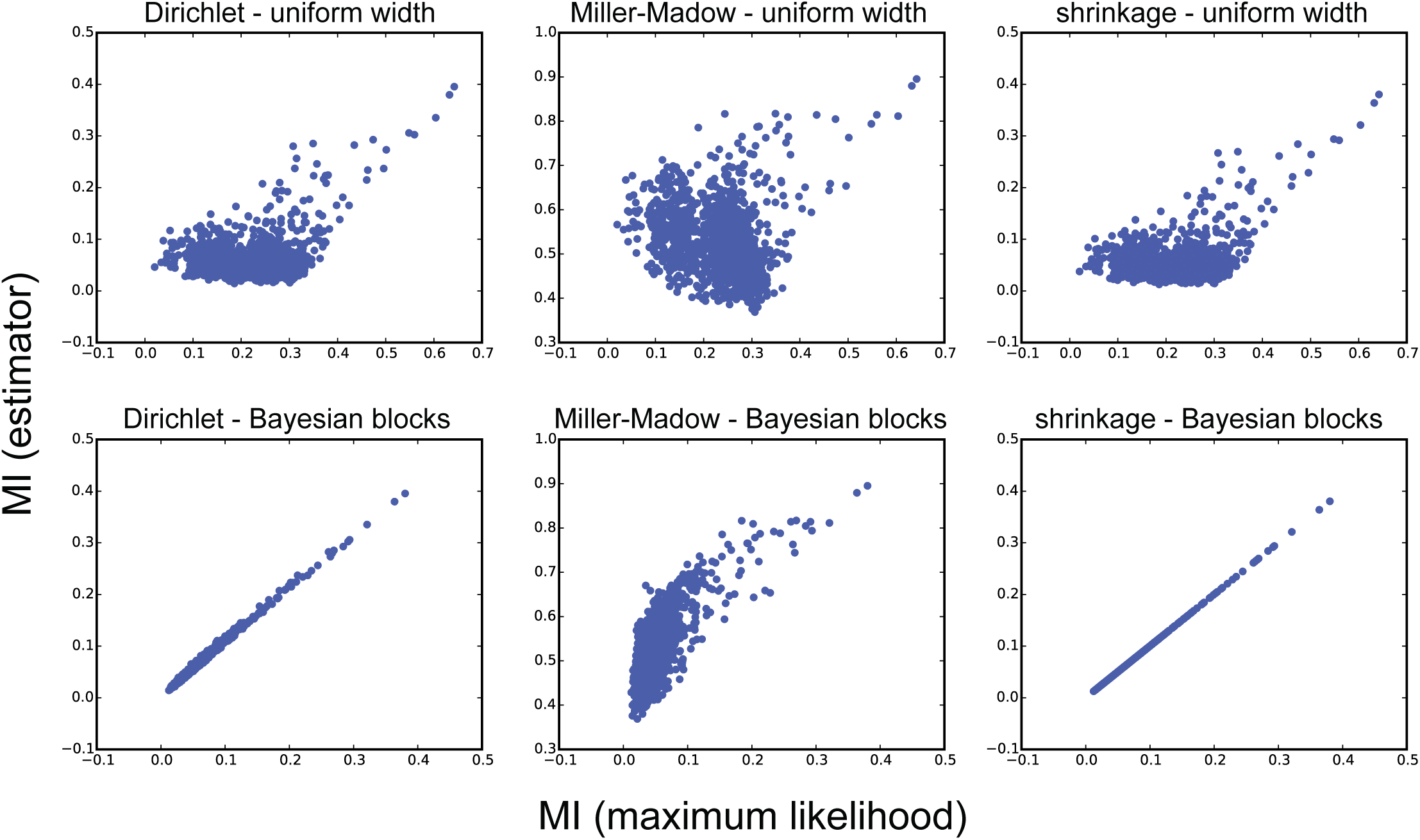
Related to Figure 5. Influence of discretization algorithm and estimator on MI rank. MI was estimated for every pair of genes in a 50-gene *in silico* network, using different combinations of discretization algorithms (uniform width or Bayesian blocks (Vanderplas et al. 2012, Scargle et al. 2013)) and MI estimators (Dirichlet, Miller-Madow, shrinkage and maximum likelihood); see *Methods* for details. Each plot shows the relative ranks of MI scores obtained using the maximum likelihood estimator (horizontal axis) versus one of the other estimators (vertical axis); the top and bottom rows show results obtained using data discretized using a uniform width or Bayesian blocks algorithm respectively. MI ranks were the most consistent when using Bayesian blocks discretization (with the exception of the Miller-Madow estimator, which is an entropy bias correction that should not be used in higher-dimensional estimation and is included here due to its frequent misuse as an MI estimator).

**Table S1:** Related to Figures 2 and S2. Topological characteristics of the 50-node and 100-node (in brackets) networks from which data were simulated by GeneNetWeaver (Schaffter et al. 2011). In all networks the vast majority of all possible node triples were either unconnected or had only one edge connecting two nodes in the triple.

| Network | Nodes | Triples | Edges | Unconnected<br>triples | One edge<br>triples |
| --- | --- | --- | --- | --- | --- |
| *E. coli* 1 | 50 (100) | 19600 (161700) | 62 (125) | 86.4% (92.8%) | 12.0% (6.8%) |
| *E. coli* 2 | 50 (100) | 19600 (161700) | 82 (119) | 82.8% (93.2%) | 14.4% (6.3%) |
| *S. cerevisiae* 1 | 50 (100) | 19600 (161700) | 77 (166) | 82.7% (90.3%) | 15.9% (9.3%) |
| *S. cerevisiae* 2 | 50 (100) | 19600 (161700) | 160 (389) | 66.6% (78.8%) | 28.2% (18.9%) |
| *S. cerevisiae* 3 | 50 (100) | 19600 (161700) | 173 (551) | 64.4% (71.6%) | 29.7% (24.1%) |

**Table S2:** Related to Figure S3 and Table S3. Joint entropy estimates, in bits, for up to four independent random variables, calculated using different estimators and uniform width discretization. Each variable is uniformly distributed over *n* = 64 bins and sampled 64^2^ times, with theoretical entropy, log_2_(*n*). The theoretical joint entropy of independent variables is the sum of their entropies. The Dirichlet estimator is given priors of 1*/n* or 1, and the shrinkage estimator is given a uniform target (as indicated in parentheses in table headings). The means of 100 repetitions are given, and the variances are all *<* 10^−4^. Only the Dirichlet and shrinkage estimators produce accurate estimates with three and four variables.

| Number of<br>variables | Theoretical<br>value | Maximum<br>likelihood | Miller-Madow | Dirichlet<br>($1/n$) | Dirichlet<br>(1) | Shrinkage<br>(uniform) |
| --- | --- | --- | --- | --- | --- | --- |
| 1 | 6 | 5.9886 | 5.9963 | 5.9892 | 5.9998 | 6.0000 |
| 2 | 12 | 11.1705 | 11.4860 | 11.2234 | 11.8303 | 11.9998 |
| 3 | 18 | 11.9844 | 12.4804 | 15.9346 | 17.9915 | 17.9999 |
| 4 | 24 | 11.9997 | 12.4804 | 23.9283 | 23.9999 | 23.9998 |

**Table S3:** Related to Figure S3 and Table S2. Estimates of the difference between joint and marginal entropies of four independent random variables. The theoretical difference for independent variables is 0. Estimates are made for three sets of variables, drawn from three different distributions (uniform, normal or exponential). The Dirichlet estimator is given priors of 1*/n* and 1, and the shrinkage estimator is given a uniform target (as indicated in parentheses in table headings). The means of 100 repetitions are given, and the variances are all < 0.2. The estimators perform differently depending on the distribution: the Dirichlet and shrinkage estimator are the most accurate when given the correct prior, but can be the least accurate when the prior is wrong.

| Distribution | Maximum<br>likelihood | Miller-Madow | Dirichlet<br>($1/n$) | Dirichlet<br>(1) | Shrinkage<br>(uniform) |
| --- | --- | --- | --- | --- | --- |
| Uniform | 11.9559 | 11.4868 | 0.0277 | 0.0428 | 0.0005 |
| Normal | 8.7172 | 8.2470 | 3.2126 | 3.1409 | 3.1389 |
| Exponential | 5.2646 | 4.8102 | 6.7636 | 6.5397 | 6.1207 |

